# Trans-synaptic assemblies link synaptic vesicles and neuroreceptors

**DOI:** 10.1101/2020.07.17.208173

**Authors:** Antonio Martinez-Sanchez, Ulrike Laugks, Zdravko Kochovski, Christos Papantoniou, Wolfgang Baumeister, Vladan Lucic

**Affiliations:** Max Planck Institute of Biochemistry, Am Klopferspitz 18, 82152 Martinsried, Germany; Institute of Neuropathology, University Medical Center Göttingen, Göttingen, Germany; Cluster of Excellence “Multiscale Bioimaging: from Molecular Machines to Networks of Excitable Cells” (MBExC), University of Göttingen, Germany

## Abstract

Synaptic transmission is characterized by fast, tightly coupled processes and complex signaling pathways that require a distinctly non-random spatial organization of their components. Nanoscale organization of synaptic proteins at glutamatergic synapses was suggested to regulate synaptic plasticity, the process underlying learning and memory. Specifically, direct colocalization of pre- and postsynaptic proteins implicated that the alignment of neurotransmitter release sites with neurotransmitter receptors enables maximal synaptic response. However, direct visualization and the mechanistic understanding of this alignment is lacking. Here we used cryo-electron tomography to visualize synaptic complexes in their native environment with the full complement of their interacting partners, synaptic vesicles and plasma membranes on 2-4 nanometer scale. The application of our recent template-free detection and classification procedure showed that tripartite trans-synaptic assemblies (subcolumns) link synaptic vesicles to postsynaptic receptors, and established that a particular displacement between directly interacting complexes characterizes subcolumns. Furthermore, we obtained de novo average structures of ionotropic glutamate receptors in their physiological composition, embedded in lipid membranes. The data presented support the hypothesis that synaptic function is carried by precisely organized trans-synaptic units. It complements superresolution findings and provides a framework for further exploration of synaptic and other large molecular assemblies that link different cells or cellular regions and may require weak or transient interactions to exert their function.

## Introduction

Most cellular processes are carried out by molecular assemblies that form functional modules and require a distinctly non-random spatial organization of their components [1, 2, 3]. Several postsynaptic, presynaptic and cell-adhesion proteins, as well as neurotransmitter release sites are organized in nanoscale domains [4, 5, 6, 7, 8, 9, 10, 11, 12, 13]. The importance of these nanodomains is exemplified by the findings that diffusion of α-amino-3-hydroxy-5-methyl-4-isoxazole propionic acid receptors (AMPARs) along the postsynaptic membrane of glutamatergic synapses can be arrested within specific nano-domains [6, 7, 14] and is required for mechanisms underlying higher cognitive processes [15, 16]. Furthermore, evidence is accumulating that the response at glutamatergic synapses depends on the exact positioning of postsynaptic receptors in respect to presynaptic release complexes, and that colocalization of nanodomains forms functional assemblies that may control synaptic strength [17, 18]. The finding that nanodomains containing some of the pre- and postsynaptic proteins directly involved in synaptic transmission colocalize across the synaptic cleft to form trans-synaptic nanoassemblies (termed synaptic nanocolumns) was a crucial step in that direction [5]. While these observations, obtained by super-resolution fluorescence microscopy, reach a precision of few tens of nanometers, direct observation of these nanoassemblies is currently lacking. Furthermore, many fundamental aspects of trans-synaptic nanoassemblies are still unresolved, such as the mechanism responsible for the trans-synaptic alignment and the existence of basic discrete units within nanocolumns.

Ionotropic glutamate receptors (iGluRs) mediate synaptic transmission at excitatory synapses [19]. Among them, AMPAR and N-methyl-d-aspartate receptors (NMDARs) are prominent components of synaptic nanodomains and are essential for long term potentiation [20]. They are composed of four pore-forming subunits, but may contain several auxiliary subunits and form larger complexes [21, 3]. While a recent series of high resolution structures of AMPARs and NMDARs provided important information about their function, these structures were obtained from reconstituted receptors that were altered in different ways to improve their expression and stability [22, 23, 24, 25, 26, 27, 28]. Therefore, there is a need to obtain structures of AMPAR and NMDAR in their native physiological conformations, subunit composition and lipid environment.

Cryo-electron tomography (cryo-ET) is uniquely suited for the comprehensive, three dimensional (3D) imaging of molecular complexes in their native context at the molecular resolution [29, 30, 31]. Cellular samples are rapidly frozen and imaged in the same vitrified, fully hydrated state [32, 33]. This is in contrast to procedures that utilize chemical fixation, dehydration and heavy-metal staining, which although essential for our current understanding of cellular ultrastructure, are known to induce membrane deformation, rearrangements and aggregation of cytosolic material, thus obscuring fine biological structures and precluding molecular interpretation [34, 35, 36, 37, 38, 39].

To overcome problems arising from the presence of many molecular species and molecular crowding, here we extended the procedure we recently developed for automated, template-free detection and unsupervised classification of heterogeneous membrane-bound complexes [40] (Figure S1). This allowed us to localize different synaptic complexes at a single nanometer precision, obtain their native average structures, explore their relation to synaptic vesicles (SVs) and synaptic plasma membranes, and resolve the trans-synaptic assemblies that they form.

## Detection of synaptic complexes in cryo-tomograms

Cryo-electron tomograms of rodent neocortical synaptosomes analyzed in this study showed smooth, continuous membranes, non-aggregated cytosolic material and well-resolved intra- and extracellular complexes, in agreement with previous cryo-ET investigations of synaptosomes and neuronal cultures [41, 42, 43, 44, 45, 39].

Cryo-ET density was traced in 3D using the discrete Morse theory segmentation and the topological persistence simplification [46, 40]. The resulting network of greyscale minima, saddle points and arcs provided an accurate representation of the proteins and lipids present at the synapse (Figure 1a, b, Supplemental video 1). By imposing geometrical constraints on the detected density, we detected membrane-bound complexes located at different synaptic layers: SV tethers (presynaptic cytosolic layer), extracellular presynaptic and extracellular postsynaptic complexes (see the Methods) (Figure 1c). We also determined centroids of spatial clusters of tethers. In this way, density tracing and detection of complexes (together comprising particle picking) yield particle sets where template-based picking bias is avoided [47].

**Figure 1:**
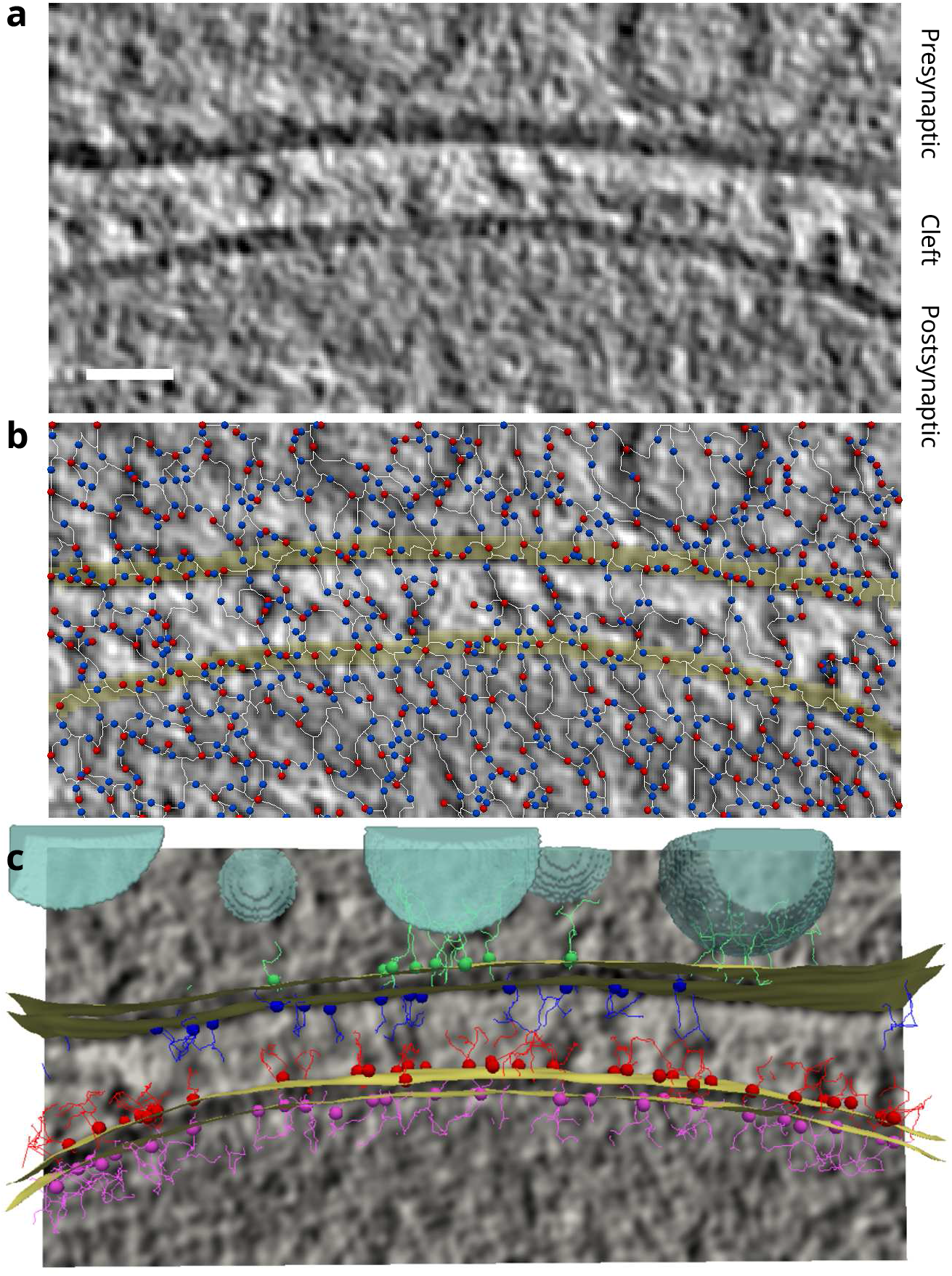
Detecting synaptic complexes. **a** Tomographic slice, 0.68 nm in thickness. **b** Tracing density using the procedure based on the discrete Morse theory. Red circles represent greyscale minima (graph vertices), the blue ones the saddle points and white lines arcs. Only for this panel the tracing was done on a single tomographic slice in 2D, all other processing was performed on tomograms, in 3D. **c** Detected particles: tethers (green), extracellular presynaptic (blue) and extracellular postsynaptic (red). Synaptic vesicles are shown in turquoise and plasma membranes in yellow color. Scale bar 20 nm.

## Classification of synaptic complexes

The particle sets obtained comprise different types of synaptic membrane-attached complexes, which reflects the inherent variability of molecular compositions and conformational states. The number and abundance of structurally similar types of complexes present at the synapse is not known a priori. We used the affinity propagation (AP) clustering in order to classify the particle sets according to their morphological features and remove false positive complexes, because AP is better suited for this task than the other commonly used methods [48, 40]. As rotational averages around vectors normal to the plasma membrane were subjected to AP clustering, the results were sensitive to the particle orientation with respect to the membrane, but not to the rotation around the membrane normal.

The first AP classification served to select extracellular particles that showed clear membrane-bound densities, yielding pre AP1 and post AP1 particle sets (Figure S2a, c). The discarded classes likely contained complexes that had weak or flexible membrane attachment domains, as well as classes composed of a mixture of heterogenous complexes. In the second AP classification round, classes lacking a well-resolved extracellular density were discarded yielding pre and post AP2 sets (Figure S2b, d). The remaining classes were merged based on visual similarity into four presynaptic (pre AP2p-s) and four postsynaptic classes (post AP2a, b, c and n), and their de novo (external reference-free) 3D averages were determined (Figures 2a, b,).

**Figure 2:**
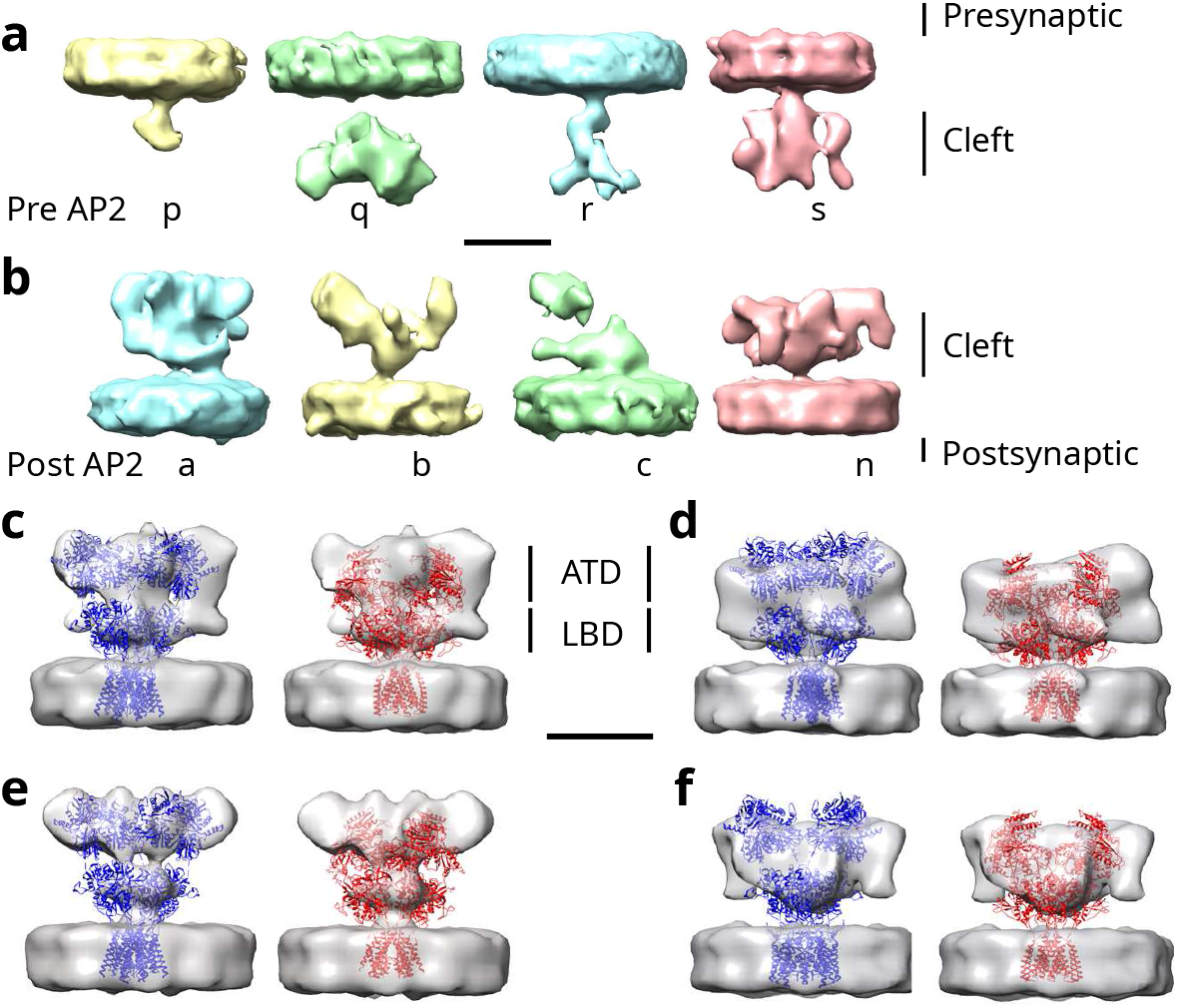
De novo average densities of extracellular presynaptic complexes. **a** Presynaptic AP2 class averages, from left to right AP2p-s. **b** Postsynaptic AP2 class averages, from left to right AP2a, b, c and n. **c** Postsynaptic AP2a class average. **d** Postsynaptic AP2n class average. **e** 3D class average with C2 symmetry derived from AP2a. **f** 3D class average with C2 symmetry derived from AP2n. **c-f** In each case the same density (grey) is shown twice: on the left atomic model of AMPAR (blue, PDB-6QKZ) is fitted in the density and on the right NMDAR model (red, PDB-6MMP). Scale bars 10 nm.

These de novo class averages showed complexes of various size and distinct morphology (Figure 2a, b). Postsynaptic AP class averages were larger in size and had different shape than the presynaptic ones even though the identical procedure was applied to pre- and postsynaptic sides, confirming the resolving power of our detection and classification procedure. Complexes present in low-copy numbers were likely discarded during the AP classification because their number was insufficient to produce well-resolved averages, or attributed to the more heterogeneous classes. Therefore, we conclude that the average structures we obtained most likely represent distinct groups of pre- and postsynaptic complexes.

## Structure determination of iGluRs

Among the four postsynaptic classes, the average AP2a and AP2n densities were well resolved (Figure 2b). Subsequent de novo 3D classification of AP2a and AP2n sets obtained without and with C2 symmetry (Figure S6a, b) yielded better resolved averages (Figures S3d, 2e, f). Because members of iGluR pore-forming subunits GluAs and GluNs are among the most highly expressed postsynaptic membrane-bound proteins [49], it is expected that they are highly prominent in AP2a and AP2n sets.

The overall size of the extracellular region of all AP2a and AP2n averages (maximal height and width > 10 nm) matched that of the available iGluR structures, and it was much larger than the extracellular region of all other currently available structures of membrane-bound postsynaptic proteins (Figure 2c-f, S3a-c). Other large complexes, such as the voltage-gated Na, Ca, and K channels, do not have extracellular amino-acid sequences that are long enough to match that of iGluRs (approximately 340 kDa). Similarly, the available structures of synaptic adhesion complexes are too small to account for the size of AP2a and AP2n class averages [50, 51]. Furthermore, our averages showed the main morphological features of iGluRs, the membrane-proximal ligand binding domains (LBD) and membrane-distal amino terminal domains (ATD) (Figure 2c-f). Finally, the averages showed an approximate two-fold symmetry around the axis perpendicular to the membrane, consistent with the architecture of iGluRs composed of four pore-forming subunits with their ATDs and LBDs organized as dimer-of-dimers. Altogether, our data argues that iGluRs are major constituents of AP2a and AP2n sets.

Despite their overall similarity, post AP2a and AP2n class averages showed important differences. The AP2a class average was taller than the AP2n, consistent with the difference between AMPAR and NMDAR type iGluRs (Figures 2c, d, S3a, b, S4a, S5a). Furthermore, the AP2a average adopted a “Y-shape”, with open membrane-distal regions, characteristic of AMPARs and showed further separated LBD and ATD domains [22]. The AP2n average had a more compact form, where the lower, membrane-proximal region was more prominent, making it similar to NMDARs [52]. Other iGluR types, kainate and delta receptors, are not expected to be prominent on the postsynaptic side of neocortical synapses [53, 54]. Therefore, AP2a and AP2n classes likely represent AMPARs and NMDARs, respectively.

Next, we fit the following three iGluR atomic models in all our de novo averages: a very recent atomic model of AMPAR that has the closest to physiological subunit composition obtained so far and two recent NMDAR models that represent the array of conformational states of NMDAR [22, 24] (Figures S4a-c, S5a-c). The height and the shape of de novo averages derived from the AP2a matched better AMPAR, while those derived from AP2n particle matched better NMDAR models.

Proceeding with the atomic initial model based 3D classification and refinement of putative AMPAR (AP2a) and NMDAR (AP2n) particles (Figures S6c, d, S1), we obtained averages that were consistent with known structures (Figures S4d, S5d, e). When the initial models were exchanged, 3D classification of each set converged to averages similar to those obtained from the same particle set and not to the exchanged initial models (Figure S7), thus further supporting the molecular assignment of AP2a and AP2n sets. Together, the data presented strongly argues that our average densities show iGluRs, and that the AP2a class is dominated by AMPARs and the AP2n by NMDARs.

There were also differences between the de novo averages and the corresponding models. Extra density protruding away from the ATD of AP2a-derived averages may be caused by binding of other proteins, because other secreted and transmembrane proteins are known to bind AMPAR [21, 55] and N-terminal interactions were shown to be critical for AMPAR anchoring at the synapse [56, 57]. Depending on the NMDAR atomic model used, extra ATD density was found towards the periphery or close to the central region (Figure S5). This argues that multiple confirmations of NMDAR are present, consistent with previous structural studies showing a substantial conformational variability of NMDARs depending on subunit composition, physiological state and buffer conditions. Also, LTD domains were positioned further away from the membrane in our averages. This may mean that imaging using Volta phase plate negatively affected fitting, or that detergents used to stabilize the transmembrane domain in single particle EM studies did not fully substitute for native lipids.

Except for early work on negatively stained samples ([58, 59]), all previously reported AMPAR and NMDAR structures were obtained from reconstituted receptors that in most cases had a non-native pore-forming and auxiliary subunit composition [22, 23, 24, 25, 26, 27, 28]. Receptors were detergent-extracted, genetically modified and artificially stabilized, leading to high resolution structures and the determination of atomic models, but left the question open to which extent they correspond to native iGluRs. This is in contrast with our samples, where iGluRs have physiological subunit compositions and are kept in their native lipid membranes together with more than 30 known auxiliary subunits and binding partners [55]. Therefore, interactions between auxiliary and pore-forming subunits, as well as between ATD domain and other synaptic proteins [56, 60] could likely explain different conformations and extra densities observed in our averages.

## Trans-synaptic colocalization and subcolumns

To investigate the trans-synaptic organization of complexes, we determined the number of colocalization events between two types of complexes residing on different synaptic layers (termed 2-colocalizations, see the Methods). Specifically, 2-colocalizations between pre and post AP1, and between pre and post AP2 sets showed the highest significance among all possible 2-colocalizations involving pre and post AP1 and AP2 complexes, tether centroids and synaptic vesicles, (Figure S8a). These were significant at colocalization distances (in cis-cleft directions) at 5, 10, and 15 nm, while all other 2-colocalizations showed significance in few cases. Also, 2-colocalizations involving tethers were generally better than than the corresponding ones involving SVs.

We proceeded to detect 3-colocalizations, that is simultaneous trans-synaptic colocalization between complexes present on all three synaptic layers. Significant 3-colocalizations were obtained between tether centroids, pre AP1 and post AP1 complexes at 5, 10 and 15 nm and between tether centroids and the pre and post AP2 complexes at 5 and 15 nm (Figure 3a). In contrast, even though the number of tether centroids and SVs was similar and their locations were correlated, SV 3-colocalizations with AP1 complexes were significant at 10 and 25 nm, while those with AP2 complexes failed to reach significance.

**Figure 3:**
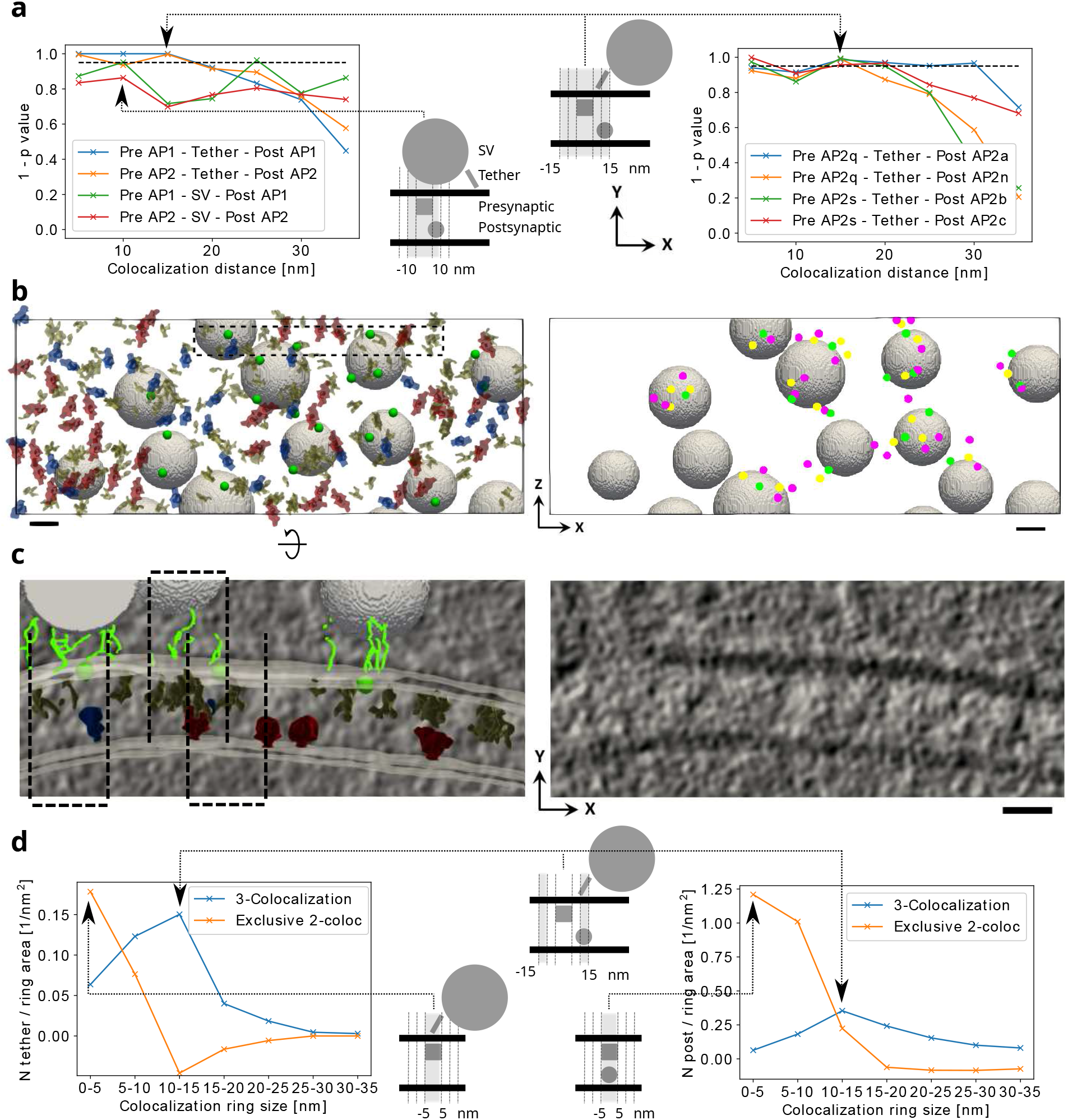
Trans-synaptic colocalization. **a** Significance of 3-colocalizations between the indicated pre-, postsynaptic complexes and tether centroids or SVs. **b** View of a synapse from the postsynaptic side where all (left) and subcolumn particles (right) are mapped at their precise locations: tether centroids (green circles), pre AP2 (yellow shapes left and yellow circles right), post AP2a (blue, left panel), post AP2n (red, left panel), all post AP2 (violet, right panel). **c** Tomographic slices corresponding to the inset shown in **b** left, viewed from the side of the synapse, with (left) and without (right) superimposed particles (green lines represent tethers, other symbols are the same as in **b**). Dashed rectangles show subcolums. Y-axes shows the trans-synaptic direction. **d** Number of tether centroids (left) and post AP2 complexes (right) in trans-synaptic rings for 3-colocalizations between tether centroids, pre AP2 and post AP2 particles (blue), and exclusive 2-colocalizations between tether centroids and pre AP2 (orange, left), and between pre and post AP2 (orange right). The data is normalized to the ring area. **a, d** (middle) Schemes in the middle illustrate single colocalization events corresponding to the most prominent data points (indicated by arrows). The shaded areas represent trans-synaptic colocalization neighborhoods (**a**) and rings (**d**). Scale bars 20 nm.

The following observations argue that 15 nm colocalization distance is the most representative of the trans-synaptic organization. The number of the tether-based 3-colocalization events normalized to the colocalization area had a maximum at 15 nm (Figure S8b). The colocalization area at 5 nm distance (78.5 nm^2^) is similar to the area of one AMPAR or NMDAR projected on the membrane and is thus too small to accommodate multiple complexes similar in size to iGluR’s within a single 3-colocalization event. Also, the number of 3-colocalization events at 5 nm was too small to support trans-synaptic organization. Given the significance of these events, they may form transient interactions. In addition, because the AP2 particle sets provide the most refined representation of the full complement of pre- and postsynaptic extracellular complexes, we will refer to the 3-colocalization events between tether centroids, pre AP2 and post AP2 complexes at 15 nm as subcolumns.

Therefore, our data strongly suggests that tripartite trans-synaptic assemblies comprising tethers, pre- and postsynaptic complexes, and subcolumns in particular, provide structural links between SVs and postsynaptic complexes and constitute a robust support for the trans-synaptic organization.

## Internal subcolumn organization

Focusing on subsets of subcolumns composed of different pre and post AP2 classes, significant results were found for the following combinations: pre AP2q - post AP2a, pre AP2q - post AP2n, pre AP2s - post AP2b and pre AP2s - post AP2c (Figures 3a, S8c). This means that among presynaptic classes, AP2q and AP2s show colocalization significance, as opposed to AP2p and AP2r. The same pattern, where AP2q and AP2s show significance, as opposed to AP2p and AP2r, was observed at 2-colocalizations between tether centroids and the pre AP2 classes (Figure S8d). Furthermore, there were clear differences between pre AP2q and AP2s complexes. 2-colocalizations tether - pre AP2s was significant for a wide range of distances, unlike tether - pre AP2q, while pre AP2q was more prominent in 3-colocalizations with iGluRs (post AP2a and AP2n) than pre AP2s. These results show that subcolumns differ in their composition, pre AP2 classes play different roles in the trans-synaptic organization and that iGluRs are prominent in subcolumns.

Next, we determined the relative particle localization within subcolumns by considering the number of particles present within concentric trans-synaptic rings (see the methods and the schemes in Figures 3d, S9c). The particle number (normalized to the ring area) within 3-colocalizations was the highest in the ring at 10-15 nm for both postsynaptic complexes and tether centroids (Figure 3d). On the contrary, a majority of tether centroids and postsynaptic complexes present exclusively in 2-colocalizations (that is in 2-but not 3-colocalizations) were located in the 0-5 nm ring, that is strongly aligned with the presynaptic complexes. Tether centroids were essentially absent from the exclusive 2-colocalization at the 10-15 nm, while the number of postsynaptic complexes present exclusively in the exclusive 2-colocalization was slightly smaller than that of postsynaptic complexes within 3-colocalization (Figure 3d). The data for tether - pre AP2q - post AP2a and tether - pre AP2q - post AP2n 3-colocalizations were very similar, except for slight shifts towards larger displacements (Figure S9c).

Therefore, our data shows that multiple forms of binary interactions exist between tethers and presynaptic, and between presynaptic and postsynaptic complexes. Furthermore, binary interactions within subcolumns are characterized by a lateral displacement between complexes (10-15 nm), as opposed to the well-aligned (0-5 nm) subcolumn - independent interactions. In addition, our data argues that the lateral positioning of tethers in the binary interaction with presynaptic complexes is strongly limiting the formation of subcolumns. This is consistent with a model where lateral and well-aligned interactions involve different proteins, but it leaves the possibility that formation of well-aligned tripartite trans-synaptic binding induces lateral shifts between its components (Figure S9e).

## Structural trans-synaptic columns

On average, there where 0.74 tether centroids, 1.36 presynaptic and 1.50 postsynaptic complexes per subcolumn. This shows that subcolumns may share components and contain multiple pre- and postsynaptic complexes, raising a possibility that subcolumns form extended structures.

Using univariate Ripley’s L function to analyze lateral spatial organization of individual particle sets, [61, 40], we found that pre AP2 and post AP2 particles did not cluster significantly (Figure S10b). As expected, tethers showed significant clustering at length scales of 5-15 nm, while tether centroids did not show significant clustering (Figure S10b).

However, taking into account only the particles located within subcolumns, both pre and post AP2 particles significantly clustered at length scales of 30-50 nm and 15-50 nm, respectively (Figure S10c). Because clustering was detected at length scales larger than the 15 nm scale used to define subcolumns, we conclude that subcolumns are organized in clusters at length scales up to at least 50 nm. Judged by the high variability and the additional maximum of the post AP2 trace at around 100 nm, it is likely that clustering range was much wider at some synapses. Therefore, we defined structural trans-synaptic columns as clusters of partially overlapping subcolumns (Figure S10a).

The mean number of structural columns per synapse was 3.6±3.4 (mean±std, N=14 synapses). On the average, 2.5±3.0 (mean±std) subcolumns formed one column, with the subcolumn area overlap of 19%±18% (mean±std). The area covered by columns per synapse was 6191±5632 nm^2^ (mean±std, three synapses containing no subcolumns excluded), or 17%±12% (mean±std) of the total active zone area. This compares with the nanocolumn size of 80 nm (corresponds to a circle of area 5027 nm^2^) and fits within the size range of other reported nanodomains (2000 - 50000 nm^2^) [6, 7, 12]. Overall, this variability shows that structural columns are not uniform and may point to a dynamic nature of their formation.

## Discussion

Synaptic transmission requires a precise coordination of pre- and postsynaptic complexes. Here we detected, visualized and analyzed at a single nanometer scale tripartite trans-synaptic assemblies comprising tethers, presynaptic and postsynaptic complexes that link SVs and postsynaptic receptors.

Several findings argue that these subcolumns constitute a structural framework underlying synaptic nanocolumns previously observed by super-resolution fluorescence microscopy [5]. (i) Subcolumns provide a structural link between SVs and iGluRs, thus providing a mechanism for nanocolumn formation [17]. (ii) Subcolumns contain tethers and iGluRs, which is consistent with nanocolumns because RIM and AMPAR subunits colocalize in nanocolumns and RIM1α is involved in tether formation [62]. (iii) Subcolumns are smaller in size than nanocolumns (diameter 30 nm vs 80 nm). (iv) Subcolumns cluster and share their components to form larger structures similar in size with nanocolumns.

The detection of assemblies comprising tethers and proteins present in the synaptic cleft supports the model whereby colocalization between neurotransmitter release and reception is mediated by intermediate complexes, rather than by neurotransmitter diffusion. Based on the radii of SVs and subcolumns, we estimate that subcolumns limit the maximal distance between release sites and receptors to 40-50 nm. Because that is still within the estimated area of maximally activated AMPARs (0.01 μm^2^, or a circle of 54 nm diameter) [17], we conclude that the formation of subcolumns enables a strong synaptic response.

The characteristic lateral displacements of components constituting subcolumns argue that subcolumn formation requires a synergistic binding of tethers, pre- and postsynaptic complexes, and suggest that indirect or bindings mediated by a diffusible protein [63, 64] may be involved. Also considering that more than the minimally required three complexes are often present within a single subcolumn and that in some cases nearby subcolumns share their constituents, multiple weak or transient interactions may contribute to the formation of structural columns.

To address the difficulties in studying synaptic complexes, which arise from their complexity and low abundance [65], we followed an approach that strikes a balance between deciphering larger molecular assemblies and maximizing resolution of individual complexes. The methods chosen allowed us to simultaneously visualize a multitude of synaptic complexes, SVs and plasma membranes at molecular resolution in their native environment. We observed iGluRs in their physiological composition within an environment containing their native membranes and interacting proteins, which controls their functional properties [21]. Because a substantial conformational variability of iGluRs was reported depending on physiological state, pore-forming and auxiliary subunit composition [66, 67, 52], AMPARs and NMDARs in our tomograms are expected to be present in a variety of conformations and with different interacting partners. Therefore, our iGluR averages emphasize physiological relevance, providing complementary information to the previously determined high resolution structures of reconstituted iGluRs.

Structural diversity of native iGluRs, together with a low number of particles, is certainly a factor that limits the resolution obtained. Further development of automated acquisition methods that addresses the difficulties in finding structures of interest should increase the number of particles and refine the identification of complexes. Previous visualizations of neurotransmitter receptors by conventional and cryo-ET of dissociated neuronal cultures [68, 45, 69] and recent advances in thinning vitrified samples [70] raise expectations that imaging synapses of intact neurons by cryo-ET will become routine. The previous work also showed the structural similarity between synaptosomes and synapses in neuronal cultures, and the importance of cryo-preparation for molecular imaging of synapses [44, 45, 39, 38], which in combination with the template-free detection and classification ([40]) is uniquely suited for precise molecular localization and identification in complex systems.

Therefore, the data we presented supports the hypothesis that synaptic function is carried by precisely organized trans-synaptic units, and provides a framework for further single nanometer scale explorations of synaptic and other molecular assemblies that require stable or transient structural interactions between multiple complexes present in different cells or cellular regions to exert their function.

## Methods

### Synaptosomal preparation

Cerebrocortical synaptosomes were extracted from 6–8 week old male Wistar rats as described previously [71, 72, 62] in accordance with the procedures accepted by the Max Planck Institute for Biochemistry. In brief, anesthetized animals were sacrificed, and the cortex was extracted and homogenized in homogenization buffer (HB; 0.32 M sucrose, 50 mM EDTA, 20 mM DTT, and one tablet of Complete mini EDTA-free protease inhibitor cocktail (Roche; 10 ml, pH 7.4) with up to seven strokes at 700 rpm in a Teflon glass homogenizer. The homogenate was centrifuged for 2 min at 2000 g, and the pellet was resuspended in HB and centrifuged for another 2 min at 2000 g. Supernatants from both centrifugations were combined and centrifuged for 12 min at 9500 g. The pellet was resuspended in HB and loaded onto a three-step Percoll gradient (3%, 10%, and 23%; Sigma-Aldrich) in HB without protease inhibitor cocktail. The gradients were spun for 6 min at 25000 g, and the material accumulated at the 10/23% interface was recovered and diluted to a final volume of 100 ml in Hepes-buffered medium (HBM; 140 mM NaCl, 5 mM KCl, 5 mM NaHCO3, 1.2 mM Na2HPO4, 1 mM MgCl2, 10 mM glucose, and 10 mM Hepes, pH 7.4). Percoll was removed by an additional washing step with HBM by centrifugation for 10 min at 22000 g, and the pellet was resuspended in HBM and immediately used in the experiments. All steps were performed at 4°C.

Synaptosomes prepared in this way are known to be capable of multiple rounds of Ca^2+^-dependent neurotransmitter release, contain a full complement of synaptic proteins and provide a robust system for cryo-ET [73, 74, 75, 49, 43, 3].

Synaptosomes were diluted to 0.7 mg/ml and incubated for 1 h at 37°C. During incubation, some synaptosomes (2 out of 17) were treated with 300 μM Glu, 300 μM Gly, and 30 μM KCl for 1 min, and some (2 out of 17) with 1 μM Phorbol-12,13-dibutyrate (PDBu) for 5 min just before vitrification. Because there were no obvious differences between treated and non-treated synaptosomes, the data was pooled together.

### Cryo-ET

For vitrification, a 3-μl drop of 10-nm colloidal gold (Sigma-Aldrich) was deposited on plasma-cleaned, holey carbon copper EM grids (Quantifoil) and allowed to dry. A 3-μl drop of synaptosomes was placed onto the grid, blotted with filter paper (GE Healthcare), and plunged into liquid ethane.

Tilt series were collected under a low dose acquisition scheme using SerialEM [76, 77] on Titan Krios [FEI] equipped with a field emission gun operated at 300 kV, with a post-GIF energy filter (Gatan) operated in the zero-loss mode and with a computerized cryostage designed to maintain the specimen temperature <-150°C. Images were recorded on a direct electron detector device (K2 Summit operated in the counting mode). Tilt series were typically recorded from −60° to 60° with a 2° angular increment. Pixel sizes was 0.34 nm and 0.42 nm at the specimen level. Volta phase-plate with nominal underfocus of 0.5 −1 μm [78] was used. The total dose was kept <100 e^−^/Å^2^. Individual frames were aligned using Motioncor2 [79]. Tilt series were aligned using gold beads as fiducial markers, and 3D reconstructions were obtained by weighted back projection (WBP) using Imod ([80]). During reconstruction, the projections were binned once and low pass filtered at the post-binning Nyquist frequency. Tomograms analyzed here showed higher contrast, thanks to the recent developments of the Volta phase plate for electron microscopes and the direct electron detectors [78, 81].

We selected for further processing 14 tomograms from four synaptosomal preparations that were of sufficient technical and biological quality. Specifically, tomograms were deemed technically acceptable if they did not contain any signs of ice crystal formation such as ice reflections or faceted membranes, and they had reasonable signal-to-noise ratio and proper tomographic alignment. Synaptosomes that showed prominent postsynaptic density indicative of the glutamatergic synapses were selected. We discarded synapses showing signs of deterioration such as elliptical small vesicles, strong endocytotic features, broken membranes, or the cleft narrower than 14 nm.

### Particle picking and classification by the affinity propagation classification

The complete image processing workflow is schematically presented in Figure S1.

Synaptic membranes and proximal SVs were segmented using an automated software approach [82], or manually by Amira software (ThermoFisher). Density tracing, particle picking and general classification was performed following the procedure described before [40]. Specifically, tomograms were smoothed by Gaussian low-pass filtering at σ=1.5 pixels, density was traced in 3D using a discrete Morse theory based software package DisPerSe [46] and simplified by the topological persistence. The persistence threshold was set so that for all synapses the density of vertices on the synaptic membranes was 0.006 vertices/nm^3^ and the number of arcs was two times higher than the number of vertices.

The Morse density tracing ensures that the vertices are located at the points of greyscale value minima (that is, density maxima), and that each arc propagates along the maximum greyscale gradient and contains a saddle point (having minima in two dimensions and a maximum in the third dimension) [83, 46]. The simplification by topological persistence reduces the influence of noise by removing spurious minima - saddle point pairs. This way, the vertices represent strong densities and the arcs the connecting structures between then. A path is composed of a chain of vertices and arcs, thus connecting more distant vertices.

Extracellular pre- and postsynaptic particles were detected from the Morse tracing results by selecting extracellular vertices that satisfy the following geometrical constraints: 8-20 nm Euclidean distance to the membrane, 20-40 nm geodesic distance (along a path) to the closest membrane vertex, sinuosity of the path 0-3 nm. The projection of selected vertices on the interface between the plasma membrane and the extracellular space was chosen as the particle center. Tethers were defined as arcs linking the SV and the presynaptic membrane vertices that had a geodesic length up to 40 nm. Exclusion (minimal interparticle) distance of 0.5 nm was imposed. Subvolumes of 64 nm in size at pixel size 0.684 nm, centered on the particle locations, were extracted for further processing, yielding 3670 extracellular presynaptic and 5688 extracellular postsynaptic particles.

For each particle, we determined the direction of the vector normal to the membrane at the exact particle position. The direction of the particle normal vectors was initially determined by calculating the direction perpendicular to the membrane at the previously determined particle positions (Supplemental video 1). We then improved the precision of the particle positions and normal vectors, by performing one round of high symmetry (C10) constrained refinement (see below). This precise determination of particle positions and normal vectors turned out to be essential for further processing.

AP classifications of pre- and postsynaptic particles followed the same procedure. Particles were rotationally averaged by computing mean greyscale values of 2 pixel-wide rings around the particle normal vectors and the resulting rotational averages were normalized to a density mean of 0 and standard deviation of 1. Clustering distance between two particles was defined as the cross-correlation coefficient between their rotational averages around vectors normal to the plasma membrane. The AP input preference parameter was −6. For the first round of AP classification, a cylindrical mask was used that included both membrane and extracellular regions (Figure S2a, c). Classes that did not show clear membrane-bound densities were discarded, leaving 8 out of 10, and 17 out of 21 pre- and postsynaptic classes, containing 1524 presynaptic (particle set pre AP1) and 2235 postsynaptic particles (post AP1 particle set). The second AP classification yielded 18 pre- and 30 postsynaptic classes. The mask was cylindrical, containing only extracellular space (Figure S2b, d). Class averages obtained from rotationaly averaged particles, together with de novo 3D refined class averages were visually inspected to select AP classes containing well-defined extracellular density, resulting in 12 pre- and 18 postsynaptic classes (pre AP2 and post AP2 particle sets, respectively). Based on their similarity, these were joined to form four pre- and four postsynaptic classes, containing 200 - 600 particles per class (pre AP2p-s and post AP2a, b, c and n particle sets) for a total of 1074 and 1100 pre- and postsynaptic particles. In this way, pre and post AP1 sets are the most comprehensive, the AP2 sets are the most representative of the full complement of extracellular complexes, while pre AP2p-s and post AP2a, b, c, n sets contain distinct classes of complexes. Exclusion distance of 5 nm (for colocalization) or 10 nm (for averaging) was imposed on each particle set before further processing.

Average 3D densities of the individual AP2 classes were determined by external reference-free (de novo), constrained 3D refinement (see below).

### 3D classification and refinement

3D classification and refinement steps were performed in Relion [84]. During the refinement, particle half-datasets were processed independently according to the “gold-standard” procedure, as implemented in Relion. The resolution was determined by Fourier shell correlation at the FSC = 0.143 criterion. Unless stated explicitly, all refinements were performed de novo, that is without the use of external references. For de novo processing, initial references were obtained by aligning all particles according to the two angles determined from normals and randomizing the third angle (around the normal direction) to remove the missing wedge. The structures shown were automatically filtered at the gold standard resolution determined by Relion. No symmetry was used unless noted otherwise.

For the constrained refinement and 3D classification, particle alignment was optimized by allowing only small changes of angles corresponding to the direction of the normal vector and small spatial displacements. The alignment around the third angle (around the normal vector) was optimized over the entire angular range, except when a high symmetry was used (C10). To implement this, we set the prior values for angles *tilt* and *psi* in Relion particle star files to the two angles defining the normals to the membrane, and specified small values (3.66) for the standard deviations of these two angles in the refine command options. In case of the constrained refinement of AP2 class averages, particles were randomly rotated around membrane normals in order to remove any possible bias arising from a previous assignment of the angle around the normals.

De novo post AP2a average contained 566 and post AP2n 532 particles. In both cases 10 nm exclusion distance was imposed to avoid double picking. De novo 3D refinements of iGluRs were performed independently on post AP2a and post AP2n sets. De novo 3D class averages were obtained from 317 (AMPAR), and 177 NMDAR particles. Initial model based 3D classification and averaging used PDB-6QKZ (AMPAR) and PDB-6MMB (NMDAR) atomic models. These models were Gaussian low-pass filtered to 6 nm. The initial-model based 3D class averages contained 257 (AMPAR) and 120 (NMDAR) particles.

To obtain a robust estimate of the receptor height in Figure S3b, density traces show the 10th highest greyscale value on each z-slice. Essentially the same results were obtained for the whole range between the 1st and 100th grayscale value.

### Colocalization

The 2-colocalization between particle sets located on different synaptic layers (presynaptic cytosolic, extracellular presynaptic and extracellular postsynaptic) at a given colocalization distance is defined as the number of particles that have at least one particle from the other particle set in their colocalization neighborhoods. The colocalization (or trans-synaptic) neighborhood of a particle is defined as the cylinder centered at the particle, its axis perpendicular to the presynaptic membrane and its base radius equal to the colocalization distance (see the schemes in Figure S8). The colocalizations were implemented by projecting all particles on the intracellular interface of the presynaptic membrane and calculating the 3D Euclidean distance between projected particles originating at different layers. The distance calculated in this way represent only the lateral (cis-synaptic) distance between particles, the distance in the trans-synaptic direction is irrelevant. All colocalizations were calcuclated for colocalization distances of 5, 10, … 35 nm.

Colocalizations between three different particle sets (3-colocalizations) were defined in the same way, except that at least one particle from each of the three different particle sets has to belong to the colocalization neighborhood (see the schemes on Figure 3a). Exclusive 2-colocalizations were defined as those 2-colocalizations that were not present in the 3-colocalizations, at the same colocalization distance. The number of particles in exclusive 2-colocalizations is obtained by subtracting the number of particles in the corresponding 3-colocalizations from the 2-colocalizations. We note that this result may be negative, as a result of a particular localization of particles within 3-colocalizations. For example, if particles from sets A and B colocalize at 5 nm distance, but a particle from set C colocalizes with the other two only at 10 nm, the number of exclusive 2-colocalizations at 5 nm is 1, but at 10 nm it is −1.

Colocalizations were also calculated for neighborhoods in the form of concentric trans-synaptic rings, both for 2- and 3-colocalizations (see schemes in Figures 3d and S9c). These were calculated by subtracting colocalization numbers between consecutive distances.

In some cases, the data was presented as the number of colocalizations at a given colocalization distance divided by the corresponding colocalization area (area of a circle of radius equal to the colocalization distance) (Figure S8b), or divided by the area of colocalization rings (Figures 3d and S9c).

Colocalization analysis involved pre and post AP1 and AP2 particle sets, AP2 classes, tethers and SVs (65 in total). We used centroids obtained by spatially clustering tethers to represent tethers (137 in total), in order to remove the influence of clustering of tethers induced by SVs on the colocalization results. Additional benefit of this approach was that the number of tether centroids was comparable to the number of SVs, which facilitated comparison between the corresponding colocalizations. The position of a SV was defined as the position of its closest voxel to the presynaptic membrane. For each 3-colocalization, the number of particles within the cylinders was calculated. To avoid double picking the exclusion distance of 5 nm was imposed on each set separately.

Statistical significance of 3-colocalizations was determined by Monte-Carlo random simulations. In order to avoid bias arising from a possible non-random distribution or real particle sets, random simulation models were created so that one particle set was kept fixed at the real particle locations and the another one (for 2-colocalizations) or the other two (for 3-colocalizations) simulated particle sets were generated by randomly distributing the same number of particles (as in the real sets) on the same synaptic geometry [61]. Each time, 200 random simulation cases were generated and the number of colocalizations was determined. This procedure was repeated twice for each 2-colocalization and three times for 3-colocalizations, each particle set was kept at the real locations once, resulting in 400 random simulations for each 2-colocalization and 600 for each 3-colocalization. The fraction of all simulation results that yielded a lower or equal number of 3-colocalization events compared to the real 3-colocalisation number represents the probability that the real colocalization is different from the random model. This probability is denoted (and shown on graphs) as 1 - p-value, where the p-value is then the probability that the null-hypothesis is correct.

Trans-synaptic subcolumns were defined as 3-colocalization events between tether centroids, pre AP2 and post AP2 particles at 15 nm colocalization distance. Structurally defined trans-synaptic columns were obtained by a simple union of partially overlapping subcolumns.

### Univariate particle distribution analysis

Univariate clustering of a given particle set was determined by Ripley’s L function, using Euclidean distance between particles, as explained before [85, 61, 40]. This function is often used for the analysis of spatial point patterns at multiple length scales because it considers the distribution of distances from a particle to multiple neighbors, thus providing a more comprehensive clustering information than the nearest neighbor and other first order functions [61, 40]. Statistical significance of particle clustering was determined by Monte-Carlo random simulations, where the same number of particles was randomly distributed on on the same synaptic geometry. Ripleys’s L was calculated for 200 simulations and the envelope within which 95% of the curves were located was then used to determine whether the distribution of the particle set was significantly different from the random distribution (at the p<0.05 significance level). A clustering is said to be significant at the range of length scales that starts at the lowest scale where Ripley’s L is significant and ends at the scale where Ripley’L reaches a maximum (but is still above the 95% significance), as customary in the field [86]. For clustering of subcolumns, the clustering range can start only at 15 nm because subcolumns are created by imposing 15 nm distance (Figure S10c).

### Statistics

Statistical significance for colocalizations and univariate Ripley’s L function was calculated using Monte Carlo random simulations, as explained above.

### Software methods

Particle picking, AP classification, spatial clustering and coloclalization analysis were performed in Python using PySeg package [40]. PySeg depends on Pyto [87], Numpy VTK [88] and *matplotlib* library [89]. For visualization, Paraview [90] and the UCSF Chimera package [91] software packages were used. All computations were done on Linux clusters at the computer center of the Max Planck Institute of Biochemistry.

### Code availability statement

The complete software, together with all dependencies, is installed as PySeg capsule on Code Ocean [92]. The latest version of the software is available on GitHub (https://github.com/anmartinezs/pyseg_system.git) and upon demand.

### Data availability statement

The following EM densities have been deposited in the EMDataBank: de novo subtomogram averages of complete post AP2a (EMD-11404) and post AP2n (EMD-11405) particle sets, de novo 3D class averages without and with C2 from AP2a (EMD-11406, EMD-11408) and AP2n (EMD-11407, EMD-11409) sets, and 3D class averages obtained using external initial models from AP2a (EMD-11410) and AP2n (EMD-11411) sets.

## Supporting information

Supplemental images

## Supplementary Information

Supplementary Information is available for this paper

## Acknowledgements

We would like to thank Florian Beck for useful discussions and Gabriela J. Greif and Martin Turk for critical reading of the manuscript. A.M.-S. was the recipient of a postdoctoral fellowship from the Séneca Foundation. This work was supported by the European Commission under grant FP7 GA ERC-2012-SyG_318987–ToPAG and by Max Planck Society.

## Author Contributions

AM-S designed and implemented the software; UL, ZK and CP acquired tomograms; AM-S and VL analyzed the data; WB provided resources and acquired funding; VL conceived, designed and supervised the research. VL wrote the manuscript; all authors edited the manuscript.

## References

[1] Alberts, B. The cell as a collection of protein machines: preparing the next generation of molecular biologists. Cell 92, 291–294 (1998).

[2] Robinson, C. V., Sali, A. & Baumeister, W. The molecular sociology of the cell. Nature 450, 973–982 (2007).

[3] Frank, R. A. W. et al. Nmda receptors are selectively partitioned into complexes and supercomplexes during synapse maturation. Nature communications 7, 11264 (2016).

[4] Perez de Arce, K. et al. Topographic mapping of the synaptic cleft into adhesive nanodomains. Neuron 88, 1165–1172 (2015).

[5] Tang, A.-H. et al. A trans-synaptic nanocolumn aligns neurotransmitter release to receptors. Nature 536, 210–214 (2016).

[6] MacGillavry, H. D., Song, Y., Raghavachari, S. & Blanpied, T. A. Nanoscale scaffolding domains within the postsynaptic density concentrate synaptic ampa receptors. Neuron 78, 615–622 (2013).

[7] Nair, D. et al. Super-resolution imaging reveals that AMPA receptors inside synapses are dynamically organized in nanodomains regulated by PSD95. J Neurosci 33, 13204–24 (2013).

[8] Kellermayer, B. et al. Differential nanoscale topography and functional role of glun2-nmda receptor subtypes at glutamatergic synapses. Neuron 100, 106–119.e7 (2018).

[9] Chamma, I. et al. Mapping the dynamics and nanoscale organization of synaptic adhesion proteins using monomeric streptavidin. Nature communications 7, 10773 (2016).

[10] Trotter, J. H. et al. Synaptic neurexin-1 assembles into dynamically regulated active zone nanoclusters. The Journal of cell biology 218, 2677–2698 (2019).

[11] Maschi, D. & Klyachko, V. A. Spatiotemporal regulation of synaptic vesicle fusion sites in central synapses. Neuron 94, 65–73.e3 (2017).

[12] Hruska, M., Henderson, N., Le Marchand, S. J., Jafri, H. & Dalva, M. B. Synaptic nanomodules underlie the organization and plasticity of spine synapses. Nature neuroscience 21, 671–682 (2018).

[13] Glebov, O. O. et al. Nanoscale structural plasticity of the active zone matrix modulates presynaptic function. Cell reports 18, 2715–2728 (2017).

[14] Choquet, D. & Triller, A. The dynamic synapse. Neuron 80, 691–703 (2013).

[15] Penn, A. C. et al. Hippocampal ltp and contextual learning require surface diffusion of ampa receptors. Nature 549, 384–388 (2017).

[16] Takeuchi, T., Duszkiewicz, A. J. & Morris, R. G. M. The synaptic plasticity and memory hypothesis: encoding, storage and persistence. Philosophical transactions of the Royal Society of London. Series B, Biological sciences 369, 20130288 (2014).

[17] Biederer, T., Kaeser, P. S. & Blanpied, T. A. Transcellular Nanoalignment of Synaptic Function. Neuron 96, 680–696 (2017).

[18] Chen, H., Tang, A.-H. & Blanpied, T. A. Subsynaptic spatial organization as a regulator of synaptic strength and plasticity. Current opinion in neurobiology 51, 147–153 (2018).

[19] Traynelis, S. F. et al. Glutamate receptor ion channels: structure, regulation, and function. Pharmacological reviews 62, 405–496 (2010).

[20] Nicoll, R. A. A brief history of long-term potentiation. Neuron 93, 281–290 (2017).

[21] Schwenk, J. et al. High-resolution proteomics unravel architecture and molecular diversity of native ampa receptor complexes. Neuron 74, 621–633 (2012).

[22] Herguedas, B. et al. Architecture of the heteromeric glua1/2 ampa receptor in complex with the auxiliary subunit tarp ?8. Science (New York, N.Y.) 364 (2019).

[23] Nakagawa, T. Structures of the ampa receptor in complex with its auxiliary subunit cornichon. Science (New York, N.Y.) 366, 1259–1263 (2019).

[24] Jalali-Yazdi, F., Chowdhury, S., Yoshioka, C. & Gouaux, E. Mechanisms for zinc and proton inhibition of the glun1/glun2a nmda receptor. Cell 175, 1520–1532.e15 (2018).

[25] Chen, S. et al. Activation and desensitization mechanism of ampa receptor-tarp complex by cryo-em. Cell 170, 1234–1246.e14 (2017).

[26] Twomey, E. C., Yelshanskaya, M. V., Grassucci, R. A., Frank, J. & Sobolevsky, A. I. Channel opening and gating mechanism in ampa-subtype glutamate receptors. Nature 549, 60–65 (2017).

[27] Lü, W., Du, J., Goehring, A. & Gouaux, E. Cryo-em structures of the triheteromeric nmda receptor and its allosteric modulation. Science (New York, N.Y.) 355 (2017).

[28] Tajima, N. et al. Activation of nmda receptors and the mechanism of inhibition by ifenprodil. Nature 534, 63–68 (2016).

[29] Lucic, V., Rigort, A. & Baumeister, W. Cryo-electron tomography: the challenge of doing structural biology in situ. J Cell Biol 202, 407–419 (2013).

[30] Beck, M. & Baumeister, W. Cryo-electron tomography: can it reveal the molecular sociology of cells in atomic detail? Trends in cell biology 26, 825–837 (2016).

[31] Oikonomou, C. M. & Jensen, G. J. Cellular electron cryotomography: Toward structural biology in situ. Annual Review of Biochemistry 86, 873–896 (2017).

[32] Taylor, K. A. & Glaeser, R. M. Electron diffraction of frozen, hydrated protein crystals. Science 186, 1036–7 (1974).

[33] Dubochet, J. et al. Cryo-electron microscopy of vitrified specimens. Q Rev Biophys 21, 129–228 (1988).

[34] Dubochet, J. & Sartori Blanc, N. The cell in absence of aggregation artifacts. Micron 32, 91–9 (2001).

[35] Murk, J. L. A. N. et al. Influence of aldehyde fixation on the morphology of endosomes and lysosomes: quantitative analysis and electron tomography. Journal of microscopy 212, 81–90 (2003).

[36] Bleck, C. K. et al. Comparison of different methods for thin section EM analysis of Mycobacterium smegmatis. J Microsc 237, 23–38 (2010).

[37] Li, Y. et al. The effects of chemical fixation on the cellular nanostructure. Exp Cell Res 358, 253–259 (2017).

[38] Liu, Y.-T., Tao, C.-L., Lau, P.-M., Zhou, Z. H. & Bi, G.-Q. Postsynaptic protein organization revealed by electron microscopy. Current opinion in structural biology 54, 152–160 (2019).

[39] Zuber, B. & Lucic, V. Molecular architecture of the presynaptic terminal. Current opinion in structural biology 54, 129–138 (2019).

[40] Martinez-Sanchez, A. et al. Template-free detection and classification of membrane-bound complexes in cryo-electron tomograms. Nature methods 17, 209–216 (2020).

[41] Zuber, B., Nikonenko, I., Klauser, P., Muller, D. & Dubochet, J. The mammalian central nervous synaptic cleft contains a high density of periodically organized complexes. Proc Natl Acad Sci U S A 102, 19192–19197 (2005).

[42] Lucic, V. et al. Multiscale imaging of neurons grown in culture: from light microscopy to cryo-electron tomography. J Struct Biol 160, 146–56 (2007).

[43] Fernández-Busnadiego, R. et al. Quantitative analysis of the native presynaptic cytomatrix by cryoelectron tomography. J Cell Biol 188, 145–56 (2010).

[44] Schrod, N. et al. Pleomorphic linkers as ubiquitous structural organizers of vesicles in axons. PloS one 13, e0197886 (2018).

[45] Tao, C.-L. et al. Differentiation and characterization of excitatory and inhibitory synapses by cryo-electron tomography and correlative microscopy. The Journal of neuroscience: the official journal of the Society for Neuroscience 38, 1493–1510 (2018).

[46] Sousbie, T. The persistent cosmic web and its filamentary structure–i. theory and implementation. Monthly Notices of the Royal Astronomical Society 414, 350–383 (2011).

[47] Henderson, R. Avoiding the pitfalls of single particle cryo-electron microscopy: Einstein from noise. Proceedings of the National Academy of Sciences of the United States of America 110, 18037–18041 (2013).

[48] Frey, B. J. & Dueck, D. Clustering by passing messages between data points. Science 315, 972–976 (2007).

[49] Cheng, D. et al. Relative and absolute quantification of postsynaptic density proteome isolated from rat forebrain and cerebellum. Mol Cell Proteomics 5, 1158–70 (2006).

[50] Yamagata, A. & Fukai, S. Structural insights into leucine-rich repeat-containing synaptic cleft molecules. Current opinion in structural biology 54, 68–77 (2019).

[51] Liu, H. Synaptic organizers: synaptic adhesion-like molecules (salms). Current opinion in structural biology 54, 59–67 (2019).

[52] Wang, J. X. & Furukawa, H. Dissecting diverse functions of nmda receptors by structural biology. Current opinion in structural biology 54, 34–42 (2019).

[53] Jaskolski, F., Coussen, F. & Mulle, C. Subcellular localization and trafficking of kainate receptors. Trends in pharmacological sciences 26, 20–26 (2005).

[54] Burada, A. P., Vinnakota, R. & Kumar, J. Cryo-em structures of the ionotropic glutamate receptor glud1 reveal a non-swapped architecture. Nature structural & molecular biology 27, 84–91 (2020).

[55] Jacobi, E. & von Engelhardt, J. Diversity in ampa receptor complexes in the brain. Current opinion in neurobiology 45, 32–38 (2017).

[56] Watson, J. F., Ho, H. & Greger, I. H. Synaptic transmission and plasticity require AMPA receptor anchoring via its N-terminal domain. Elife 6 (2017).

[57] Díaz-Alonso, J. et al. Subunit-specific role for the amino-terminal domain of ampa receptors in synaptic targeting. Proceedings of the National Academy of Sciences of the United States of America 114, 7136–7141 (2017).

[58] Nakagawa, T., Cheng, Y., Ramm, E., Sheng, M. & Walz, T. Structure and different conformational states of native AMPA receptor complexes. Nature 433, 545–9 (2005).

[59] Nakagawa, T., Cheng, Y., Sheng, M. & Walz, T. Three-dimensional structure of an AMPA receptor without associated stargazin/TARP proteins. Biol Chem 387, 179–87 (2006).

[60] Greger, I. H., Watson, J. F. & Cull-Candy, S. G. Structural and Functional Architecture of AMPA-Type Glutamate Receptors and Their Auxiliary Proteins. Neuron 94, 713–730 (2017).

[61] Wiegand, T. & Moloney, K. A. Rings, circles, and null-models for point pattern analysis in ecology. Oikos 104, 209–229 (2004).

[62] Fernández-Busnadiego, R. et al. Cryo-electron tomography reveals a critical role of rim1α in synaptic vesicle tethering. J Cell Biol 201, 725–740 (2013).

[63] Uemura, T. et al. Trans-synaptic interaction of glurdelta2 and neurexin through cbln1 mediates synapse formation in the cerebellum. Cell 141, 1068–1079 (2010).

[64] Matsuda, K. et al. Cbln1 is a ligand for an orphan glutamate receptor delta2, a bidirectional synapse organizer. Science (New York, N.Y.) 328, 363–368 (2010).

[65] Frank, R. A. & Grant, S. G. Supramolecular organization of nmda receptors and the postsynaptic density. Current opinion in neurobiology 45, 139–147 (2017).

[66] Chen, S. & Gouaux, E. Structure and mechanism of ampa receptor - auxiliary protein complexes. Current opinion in structural biology 54, 104–111 (2019).

[67] Kamalova, A. & Nakagawa, T. Ampa receptor structure and auxiliary subunits. The Journal of physiology (2020).

[68] Chen, X. et al. Organization of the core structure of the postsynaptic density. Proc Natl Acad Sci U S A 105, 4453–8 (2008).

[69] Liu, Y.-T. et al. Mesophasic organization of gabaa receptors in hippocampal inhibitory synapse. bioRxiv (2020). https://www.biorxiv.org/content/early/2020/01/06/2020.01.06.895425.full.pdf.

[70] Schaffer, M. et al. A cryo-fib lift-out technique enables molecular-resolution cryo-et within native caenorhabditis elegans tissue. Nature methods 16, 757–762 (2019).

[71] Dunkley, P. R. et al. A rapid percoll gradient procedure for isolation of synaptosomes directly from an s1 fraction: homogeneity and morphology of subcellular fractions. Brain Res 441, 59–71 (1988).

[72] Godino, M. d. C., Torres, M. & Sánchez-Prieto, J. Cb1 receptors diminish both ca(2+) influx and glutamate release through two different mechanisms active in distinct populations of cerebrocortical nerve terminals. J Neurochem 101, 1471–1482 (2007).

[73] Nicholls, D. G. & Sihra, T. S. Synaptosomes possess an exocytotic pool of glutamate. Nature 321, 772–3 (1986).

[74] Whittaker, V. P. Thirty years of synaptosome research. J Neurocytol 22, 735–742 (1993).

[75] Walikonis, R. S. et al. Identification of proteins in the postsynaptic density fraction by mass spectrometry. J Neurosci 20, 4069–80 (2000).

[76] Koster, A. J. et al. Perspectives of molecular and cellular electron tomography. J Struct Biol 120, 276–308 (1997).

[77] Mastronarde, D. N. Automated electron microscope tomography using robust prediction of specimen movements. J Struct Biol 152, 36–51 (2005).

[78] Danev, R., Buijsse, B., Khoshouei, M., Plitzko, J. M. & Baumeister, W. Volta potential phase plate for in-focus phase contrast transmission electron microscopy. Proc Natl Acad Sci USA 111, 15635–15640 (2014).

[79] Zheng, S. Q. et al. Motioncor2: anisotropic correction of beam-induced motion for improved cryo-electron microscopy. Nature methods 14, 331–332 (2017).

[80] Kremer, J. R., Mastronarde, D. N. & McIntosh, J. R. Computer visualization of three-dimensional image data using imod. J Struct Biol 116, 71–76 (1996).

[81] McMullan, G., Faruqi, A. & Henderson, R. Direct electron detectors. In Methods in enzymology, vol. 579, 1–17 (Elsevier, 2016).

[82] Martinez-Sanchez, A., Garcia, I., Asano, S., Lucic, V. & Fernandez, J.-J. Robust membrane detection based on tensor voting for electron tomography. J Struct Biol 186, 49–61 (2014).

[83] Forman, R. A user’s guide to discrete morse theory. Seminaire Lotharingien de Combinatoire 48, 35pp (2002).

[84] Bharat, T. A. & Scheres, S. H. Resolving macromolecular structures from electron cryo-tomography data using subtomogram averaging in relion. Nature protocols 11, 2054 (2016).

[85] Ripley, B. D. Spatial statistics (Willey-Interscience, 1981).

[86] Kiskowski, M. A., Hancock, J. F. & Kenworthy, A. K. On the use of ripley’s k-function and its derivatives to analyze domain size. Biophysical journal 97, 1095–1103 (2009).

[87] Lucic, V., Fernández-Busnadiego, R., Laugks, U. & Baumeister, W. Hierarchical detection and analysis of macromolecular complexes in cryo-electron tomograms using pyto software. Journal of structural biology 196, 503–514 (2016).

[88] Schroeder, W. J., Lorensen, B. & Martin, K. The visualization toolkit: an object-oriented approach to 3D graphics (Kitware, 2004).

[89] Hunter, J. D. Matplotlib: A 2d graphics environment. Comput. Sci. Eng. 9, 90–95 (2007).

[90] Ayachit, U. The paraview guide: a parallel visualization application (2015).

[91] Pettersen, E. F. et al. Ucsf chimera, a visualization system for exploratory research and analysis. J Comput Chem 25, 1605–12 (2004).

[92] Martinez-Sanchez, A. & Vladan, L. Pyseg: Template-free detection and classification for cryo-et. Code ocean https://doi.org/10.24433/CO.0526052.v1 (2019).

