## Supplemental images for "Trans-synaptic assemblies link synaptic vesicles and neuroreceptors"

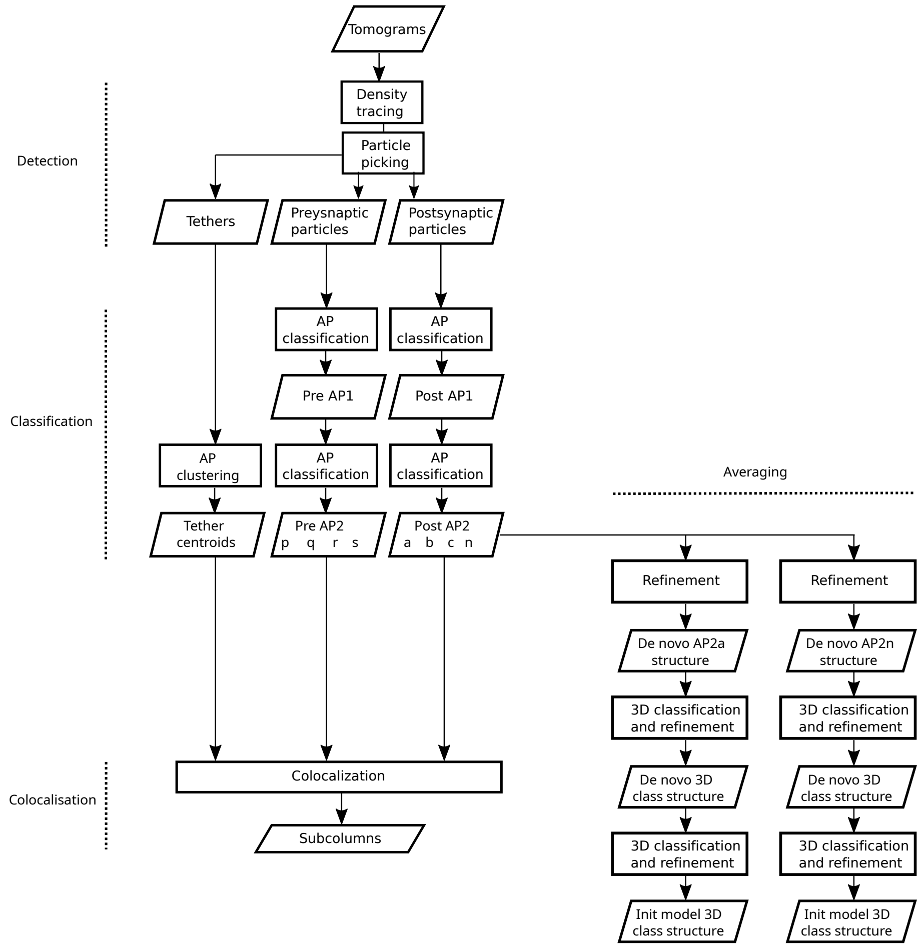


Figure S1: Processing scheme. Data is shown as parallelograms and the processing steps as rectangles.

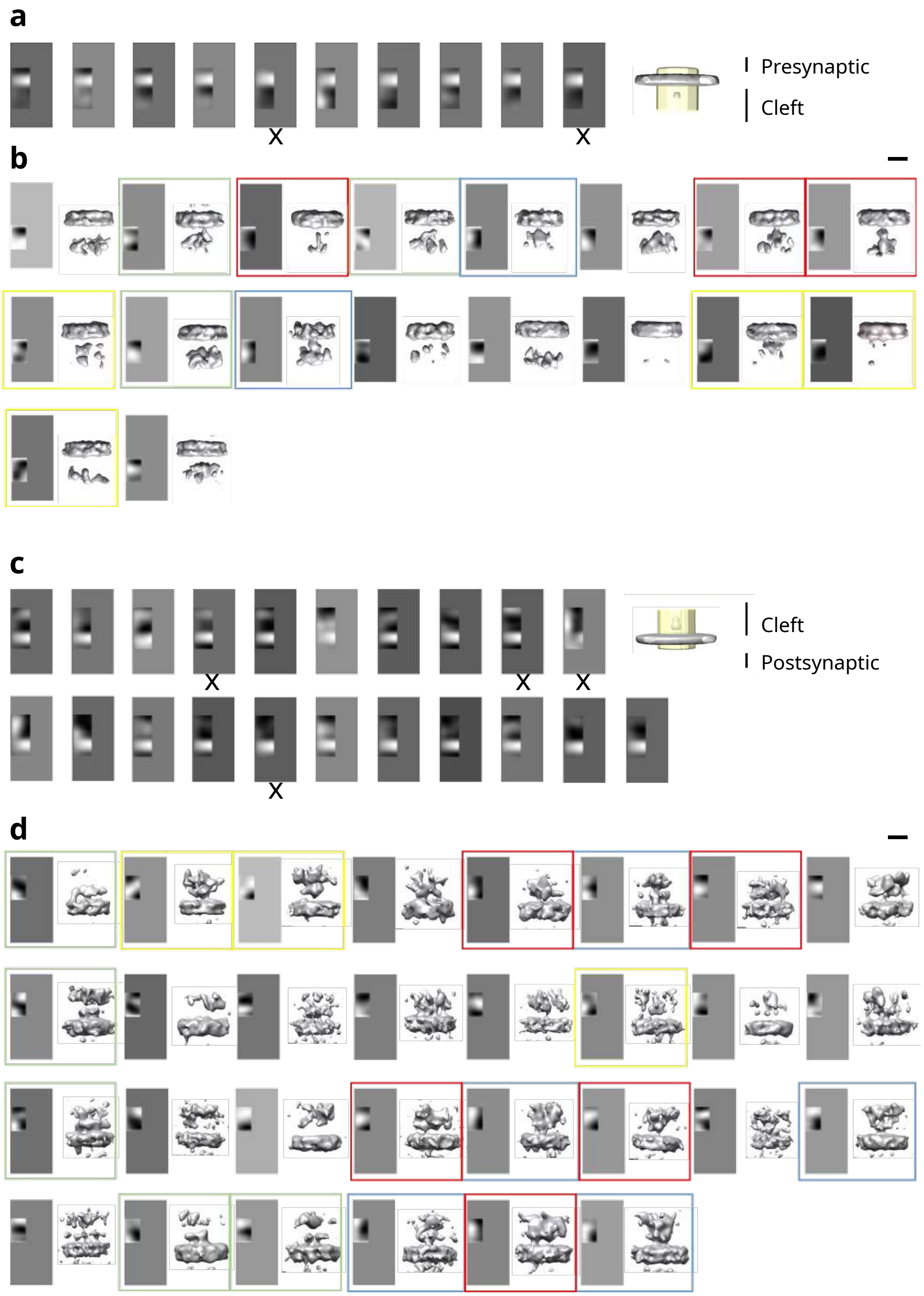


Figure S2: AP classification of pre- and postsynaptic particles. **a** First round of presynaptic AP classification, AP class averages (left), mask (transparent yellow, right). **b** Second round presynaptic AP class averages and the corresponding 3D averages. **c** First round of postsynaptic AP classification, AP class averages (left), mask (transparent yellow, right). **d** Second round postsynaptic AP class averages and the corresponding 3D averages. **a** and **c** Discarded classes are denoted by "x" below the class. AP class averages are shown as rotation averages around the normal vector. **b** and **d** Selected classes are boxed. Classes boxed by the same color are joined together to form classes shown in Figure 2a and b (colors match). Scale bars 10 nm.

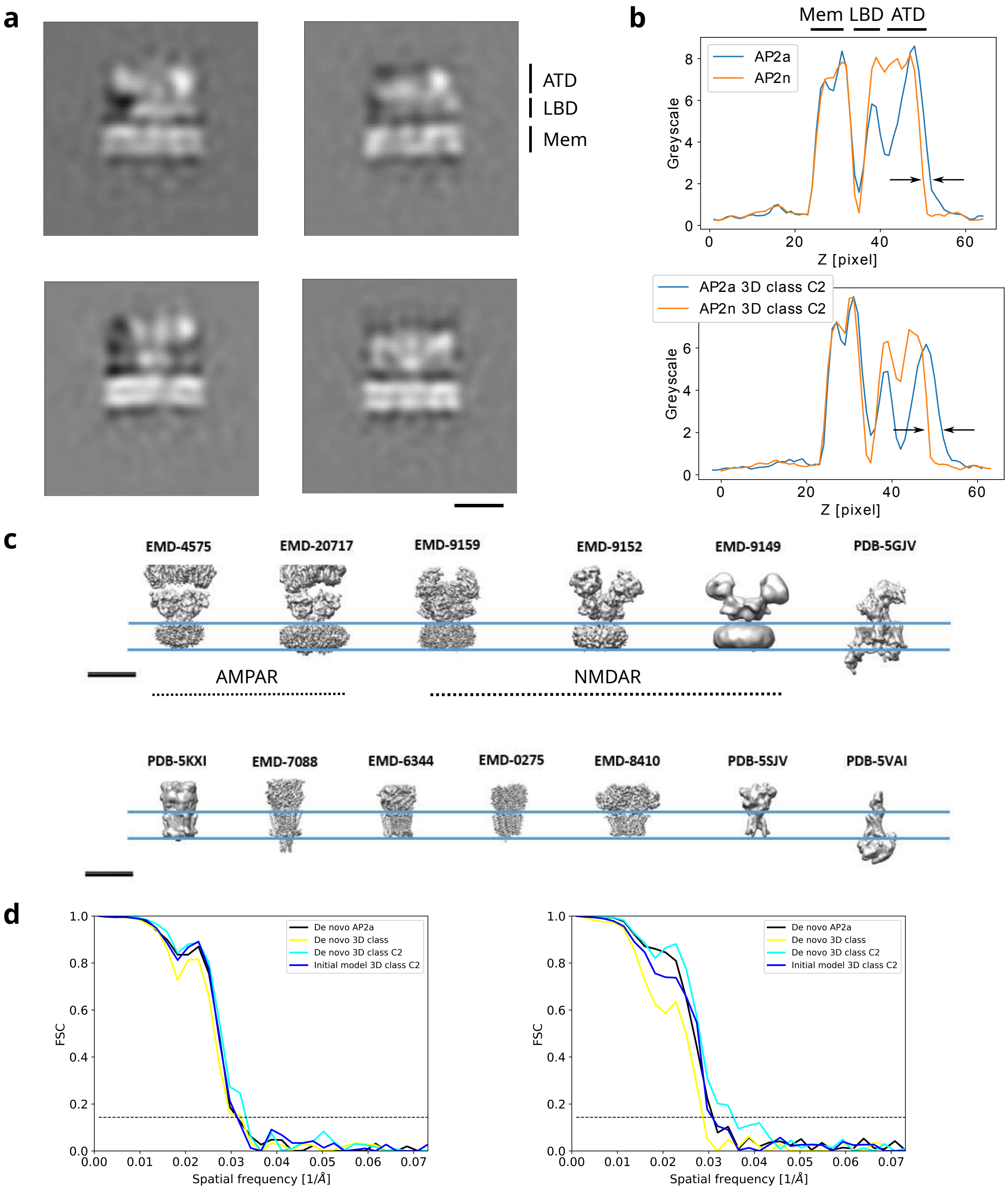


Figure S3: Averaging and identification of post AP2a and AP2n particle sets. **a** Tomographic slices of de novo averages showing the largest extent in the direction perpendicular to the membrane: AP2a (top left), AP2n (top right), 3D class with C2 from AP2a (bottom left), 3D class from AP2n (bottom right). ATD denotes amino terminal domain, LBD ligand binding domain and Mem plasma membrane. Slice thickness 0.68 nm. **b** Greyscale density traces of averages shown in panel **a**. Arrows indicate the height difference between averages. **c** Currently available structures of postsynaptic complexes. AMPAR and NMDAR denote different structures obtained from these complexes. **d** Gold-standard FSC curves for de novo and initial model based averages obtained from AP2a (left) and AP2n sets (right). ATD denotes the position of amino terminal and LBD of ligand binding domains. Scale bars 10 nm.

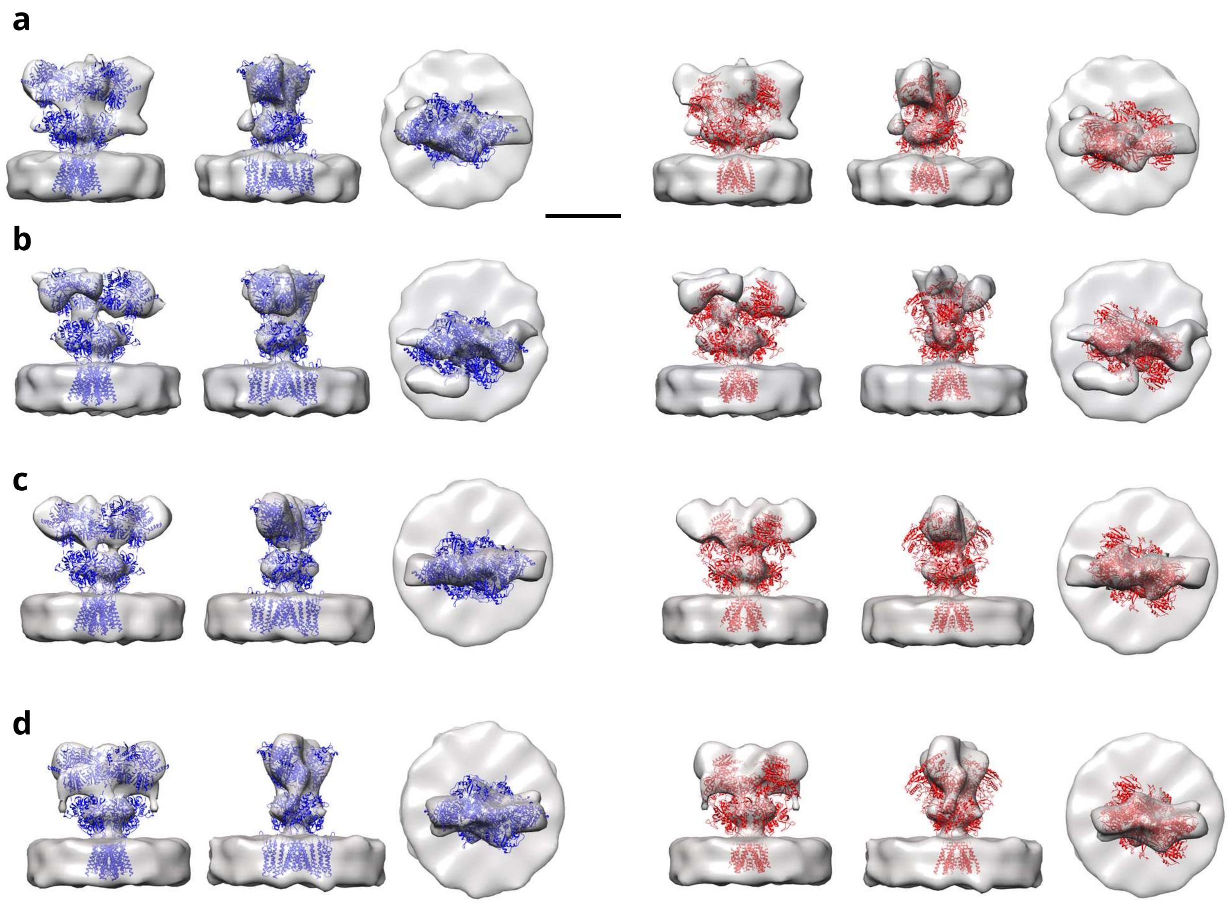


Figure S4: Subtomogram averages obtained from post AP2a set and fitting atomic models. **a**. De novo AP2a class average. **b** De novo 3D class average. without symmetry. **c** De novo 3D class averages with C2 symmetry. **d** Average obtained starting from an external initial model. All averages are shown as isosurfaces viewed from three orthogonal directions. Atomic models fit are: AMPAR (PDB-6QkZ, blue) on the left and NMDAR (PDB-6MMP, red) on the right. Scale bar 10 nm.

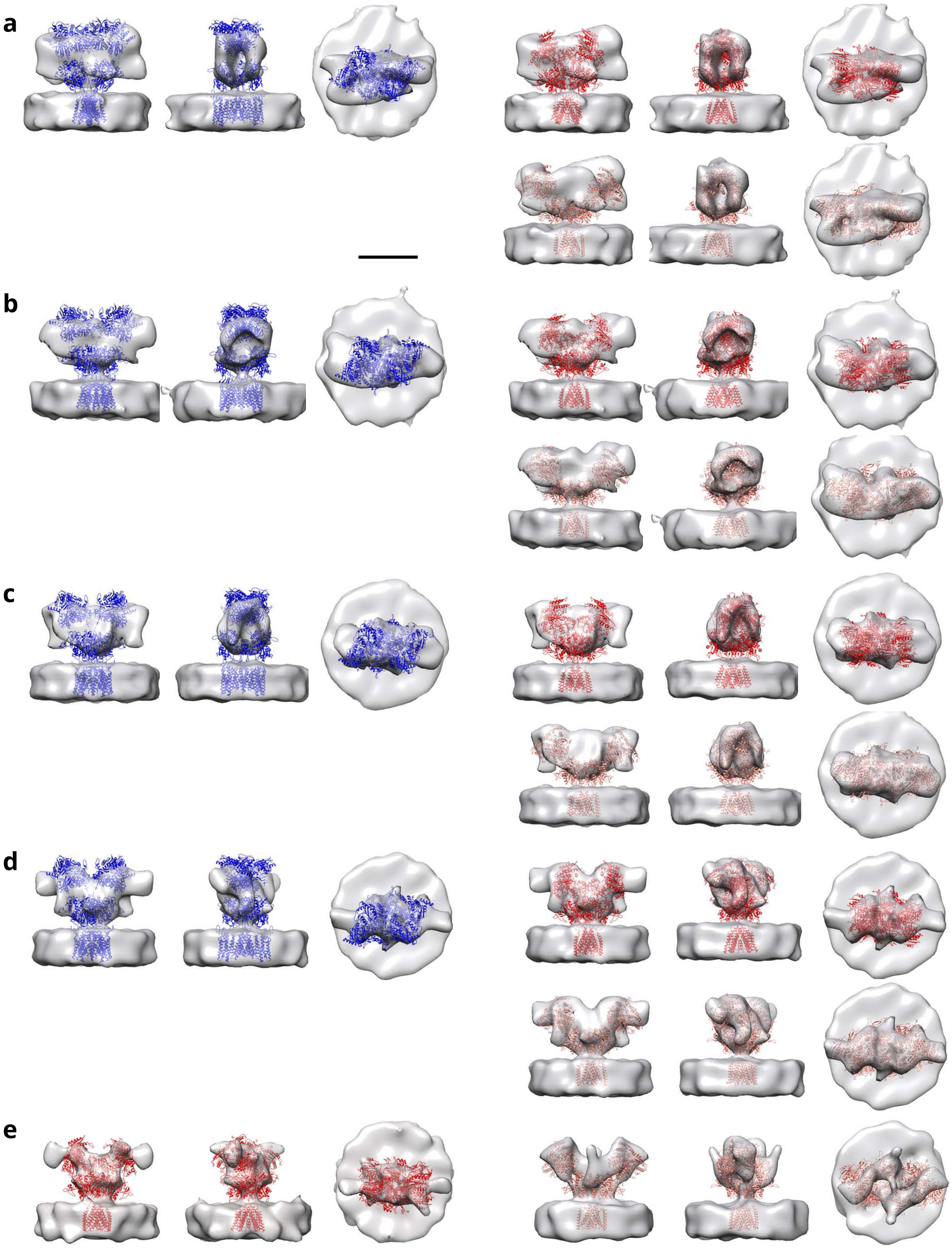
a
b
c
d
e

Figure S5: Subtomogram averages obtained from post AP2n set and fitting atomic models. **a** De novo AP2n class average. **b** De novo 3D class average, no symmetry. **c** De novo 3D class average, C2 symmetry. **d** Average obtained using an external initial model. **e** Two classes obtained from average in D. All averages are shown as isosurfaces. Atomic models: AMPAR (PDB-6QkZ, blue) on the left, NMDAR (PDB-6MMP, red) on the right top and NMDAR (PDB-6MMB, red) on the right bottom, except in **e** NMDAR (PDB-6MMP) left top and NMDAR (PDB-6MMB) right. Scale bar 10 nm.

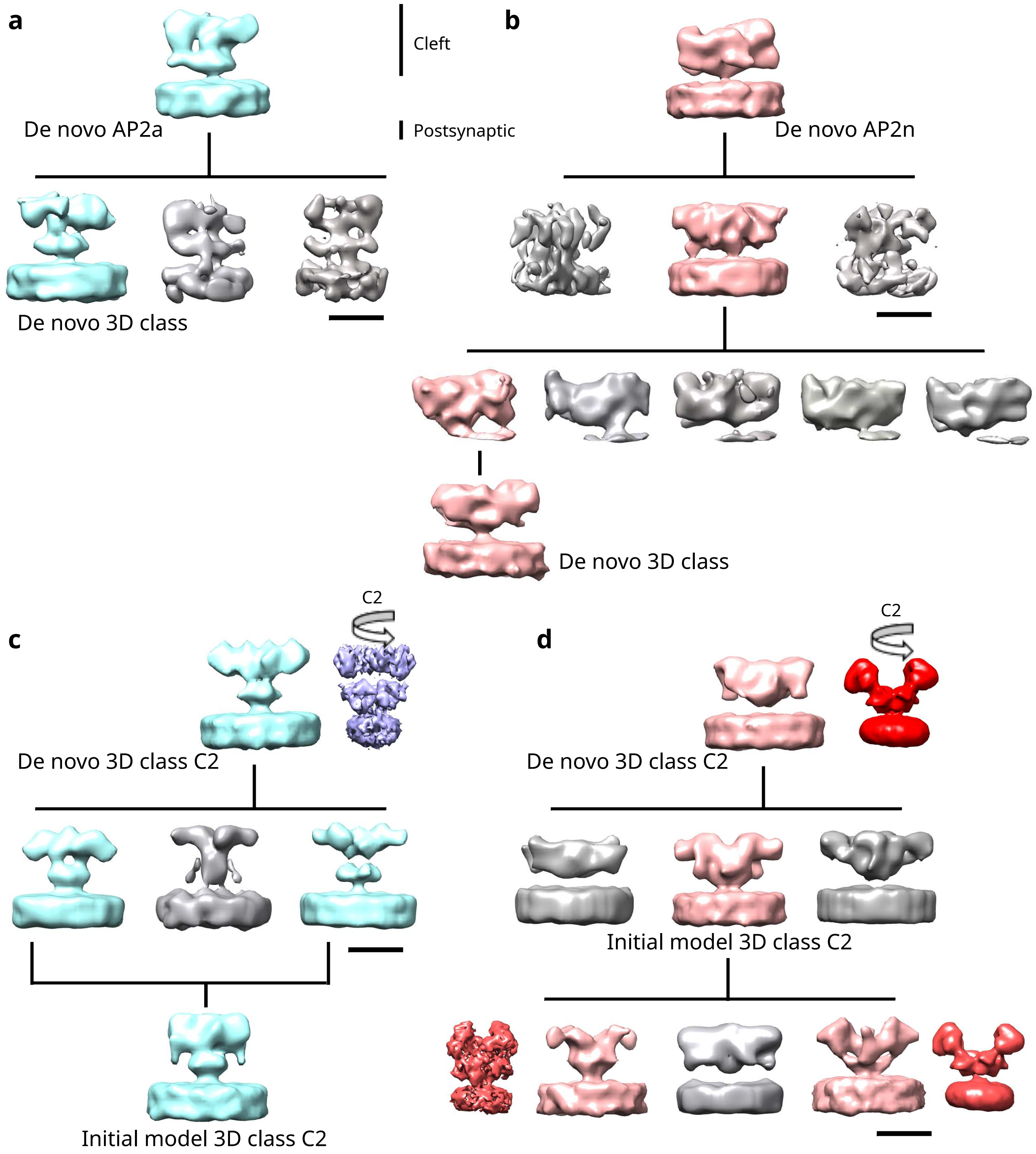


Figure S6: 3D classification of iGluRs. **a** De novo 3D classification of AP2a particles. **b** De novo 3D classification of AP2n particles. **c** 3D classification of de novo AP2a 3D class average using AMPAR atomic model (PDB-6QKZ) as initial reference. **d** 3D classification of de novo AP2n 3D class average using NMDAR atomic model (PDB-6MMB) as initial reference. The last 3D classification was done using PDB-6MMP (left) and PDB-6MMB (right) as initial references. Averages shown in color were deemed sufficiently good for further processing or final results. The labeled densities are also shown in Figures S4 and S5 and their FSC curves in Figure S3d. Scale bars 10 nm.

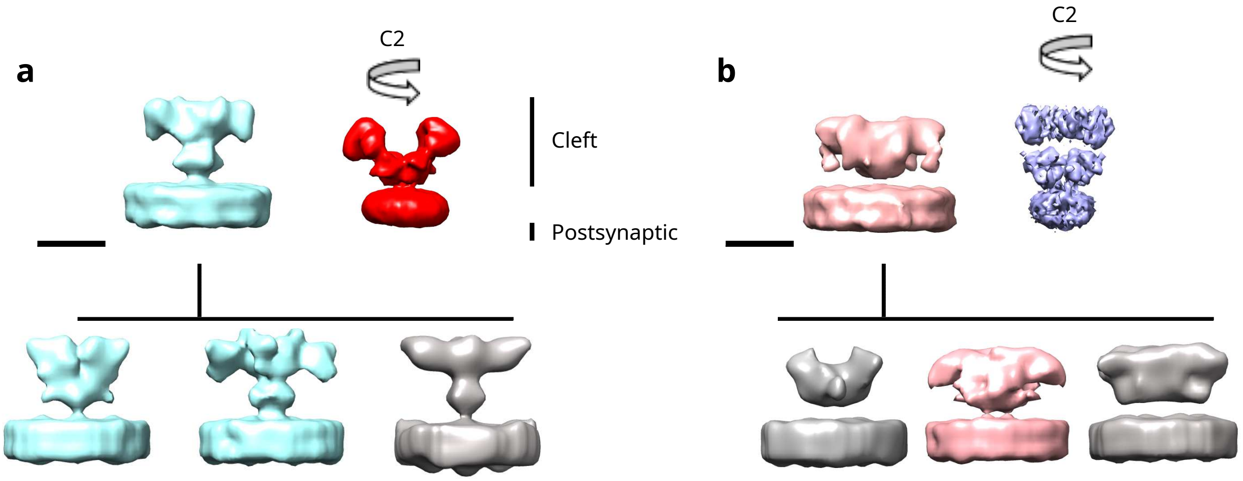


Figure S7: 3D classification of iGluRs using the exchanged initial models. **a** 3D classification of de novo AP2a 3D class average using NMDAR atomic model (PDB-6MMB) as initial reference. **b** 3D classification of de novo AP2n 3D class average using AMPAR atomic model (PDB-6QKZ) as initial reference. Averages shown in color in the lower row were deemed sufficiently good for further processing or final results. Scale bars 10 nm.

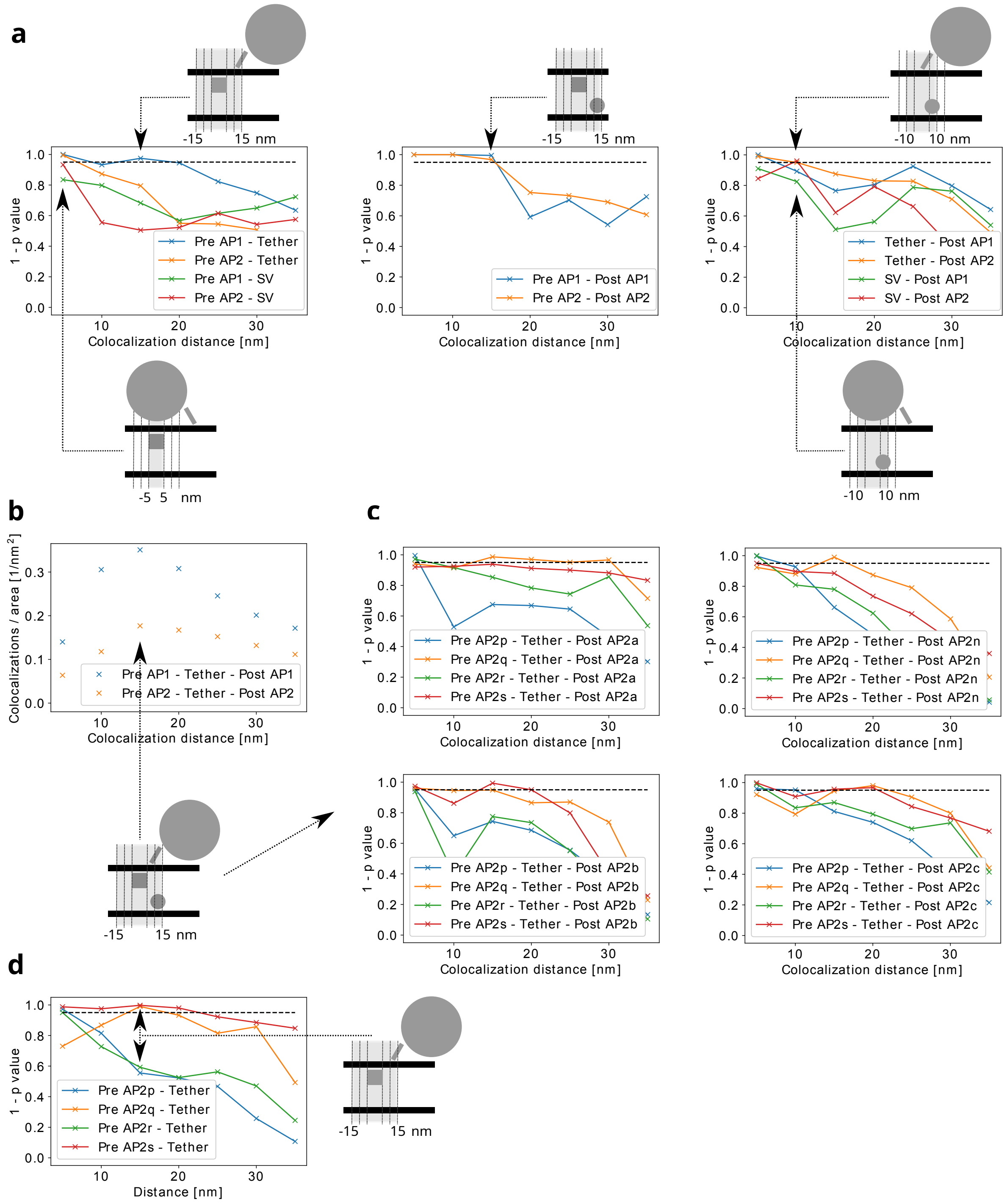


Figure S8: Trans-synaptic colocalization. **a** Significance of 2-colocalizations between the indicated pre, post-synaptic complexes and tether centroids or SVs. **b** The number of 3-colocalization events between tether centroids and the indicated pre and postsynaptic complexes. **c** Significance of 3-colocalization between the indicated tether centroids, and pre and postsynaptic AP2 classes. **d** Significance of 2-colocalizations between tether centroids and presynaptic AP2 classes. Schemes illustrate single 2-colocalization events corresponding to the most prominent data points (indicated by arrows). The shaded areas represent trans-synaptic colocalization neighborhoods.

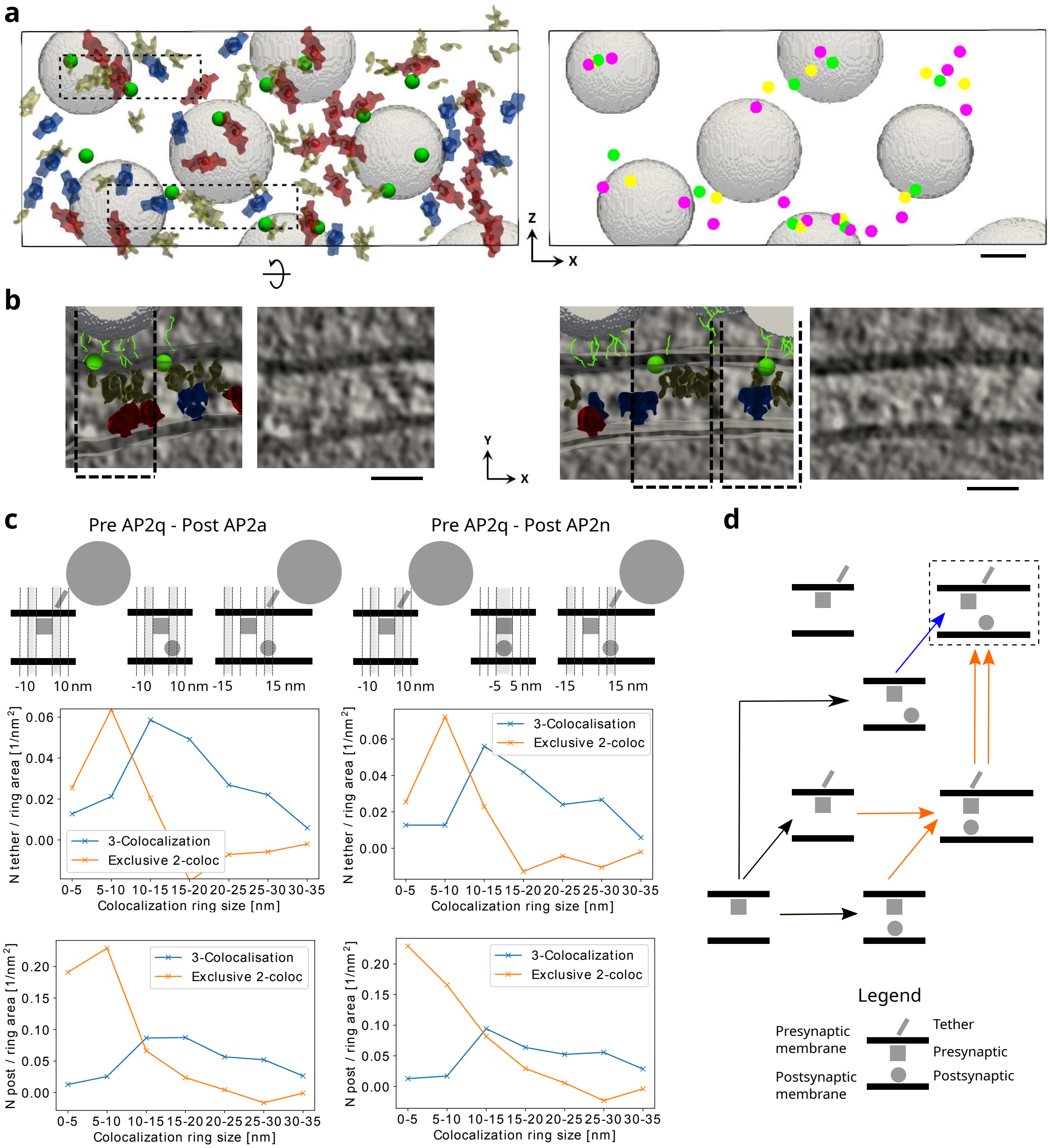

a
Z
X
b
Y
X
c
Pre AP2q - Post AP2a
Pre AP2q - Post AP2n
-10 10 nm
-10 10 nm -15 15 nm
-10 10 nm
-5 5 nm -15 15 nm
N tether / ring area [1/nm²]
3-Colocalization
Exclusive 2-coloc
3-Colocalisation
Exclusive 2-coloc
Colocalization ring size [nm]
0-5 5-10 10-15 15-20 20-25 25-30 30-35
N post / ring area [1/nm²]
d
Legend
Presynaptic membrane
Postsynaptic membrane
Tether
Presynaptic
Postsynaptic

Figure S9: Trans-synaptic subcolumns. **a** View of a synapse from the postsynaptic side where particles are mapped at their precise locations: tether centroids (green circles), pre AP2 (yellow shapes left and yellow circles right), post AP2a (blue, left panel), post AP2n (red, left panel), all post AP2 (violet, right panel). **b** Tomographic slices corresponding to insets shown in **a** left, viewed from the synapse side, with (left) and without (right) superimposed particles (green lines represent tethers, other symbols are the same as in **a**). Dashed rectangles show subcolumns. Y-axes shows the trans-synaptic direction. **c** Number of tether centroids (top row graphs) and post complexes (lower row graphs) in trans-synaptic rings for colocalizations between tether centroids, pre AP2q and post AP2a coplexes (left column), and for colocalizations between tether centroids, pre AP2q and post AP2n coplexes (right column). 3-colocalizations are shown in blue and exclusive 2-colocalizations in orange. The data is normalized to the ring area. Schemes illustrate the most prominent relative positioning of complexes. The shaded areas represent trans-synaptic colocalization rings. **d** Reaction schemes for the most likely subcolumn formation models where: lateral and well-aligned interactions have different molecular composition (blue), and well-aligned tripartite bindings induce lateral shifts (orange). Black arrows indicate model-independent transitions. Double arrow indicate a fast transition. Dashed box surrounds a subcolumn. Scale bars 20 nm.

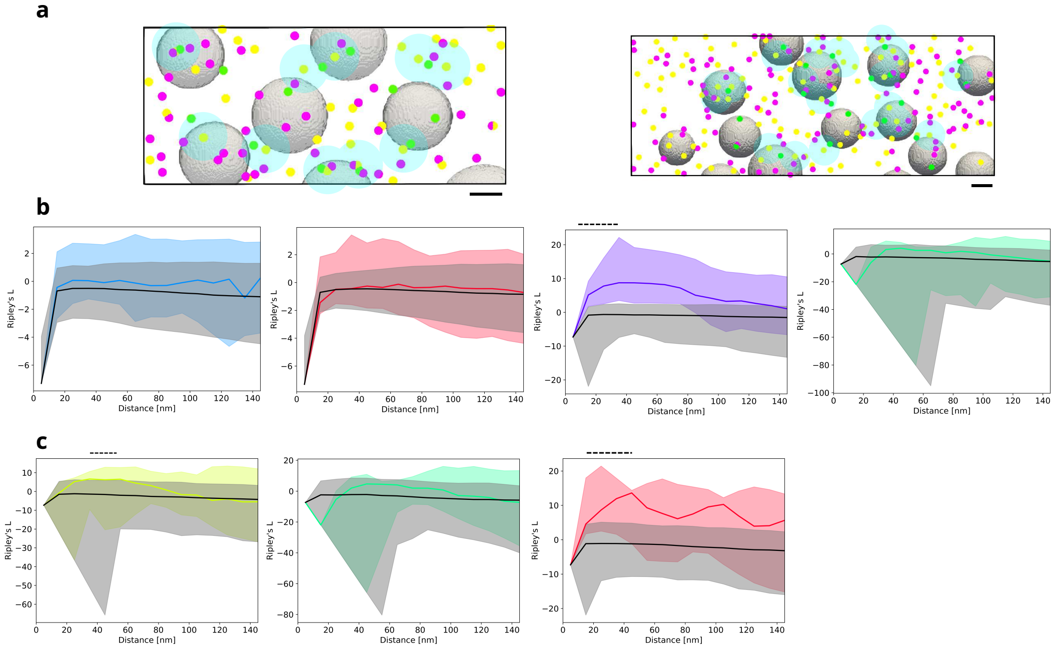


Figure S10: Structural trans-synaptic columns. **a** Examples of columns (transparent blue) superimposed on mapped particles: tether centroids (green), pre AP2 (yellow) and post AP2 complexes (violet). Scale bar 20 nm. **b** Univariate clustering of particles, from left to right: pre AP2, post AP2, tethers, tether centroids. **c** Univariate clustering of particles within subcolumns, from left to right: pre AP2, tether centroids, post AP2. Dashed black lines indicate the clustering length range. **b** and **c**, Ripley's L function is shown for real complexes (in color) and randomly positioned particles (gray). Solid lines show means, color shaded areas the variation for individual synapses and gray shaded areas the 95% confidence limits.

Video 1: Detection of synaptic complexes. This video first shows tomographic slices of a synapse (presynaptic terminal on top, synaptic cleft in the middle, postsynaptic terminal on the bottom), followed by Morse density tracing where vertices are shown as small spheres and arcs as lines: presynaptic cytoplasmic (brown), presynaptic membrane (light blue), cleft (red), postsynaptic membrane (dark blue) and postsynaptic cytosolic (grey). Next, presynaptic (blue) and postsynaptic particles (red) extracted from the tracing (small spheres) are shown together with their normal vectors (arrows). Finally, the average structures of complexes are mapped at their locations: all four pre AP2 classes (dark yellow), and 3D class averages of putative AMPAR (blue) and NMDAR (red). Scale bar 20 nm.
